# Lysine-rich regions located in the C-terminal domain of DNA Topoisomerase 2-alpha act as polyphosphoinositide interaction sites and nucleolar localisation signals

**DOI:** 10.1101/2025.01.29.635480

**Authors:** Vandana V. Ardawatia, Rhîan G. Jacobsen, Diana C. Turcu, Sandra Ninzima, Niclas P. Decker, Amanda J. Edson, Aurélia E. Lewis

**Author notes:** should be considered joint first author (Vandana V. Ardawatia, Rhîan G. Jacobsen).

## Abstract

Polyphosphoinositides (PPIn) are lipid signalling molecules that regulate essential cellular processes in eukaryotic cells by regulating effector proteins not only in the cytoplasm, but also in the nucleus. To decipher PPIn nuclear functions, we have previously established quantitative proteomic methods in combination with PPIn pull-down to identify nuclear PPIn-binding proteins. We focused our analyses on one of these proteins, DNA Topoisomerase 2α (TOP2A), an enzyme known to resolve DNA topological problems occurring during replication, transcription and mitosis. In this study, we validated its direct interaction with PPIn and identified the binding site consisting of two lysine-rich polybasic regions (PBR) located within its C-terminal domain (CTD) at residues 1228-1237 and 1259-1276. Overexpression studies of the WT and PBR deletion mutants showed that these two PBR were required for TOP2A nucleolar localisation. TOP2A is localised to different sub-nuclear sites where PPIn have also been detected. We hence investigated the localisation of TOP2A by immunofluorescence microscopy and cell fractionation in relation to PPIn. We show that TOP2A colocalised with the PPIn phosphatidylinositol(4,5)bisphosphate (PtdIns(4,5)*P*_2_) in nuclear speckles of transcriptionally arrested HeLa cells. Inhibition of type II phosphatidylinositol-5-phosphate 4-kinase, which generates PtdIns(4,5)*P*_2_ from PtdIns5*P*, led to an increase of TOP2A levels associated with the chromatin. Our results demonstrated TOP2A as a PPIn effector protein and a role for nuclear PPIn in regulating the subnuclear dynamic of TOP2A.

## Introduction

The nucleus is packed with nucleic acids and proteins but contains also lipids. Lipids provide structural support for the nuclear envelope and are, in addition, found within the confines of the nucleus (Samardak et al., 2024), as components of nuclear lipid droplets (Layerenza et al., 2013; McPhee et al., 2022; Fujimoto, 2024), associated with the chromatin (Albi et al., 2003; Hunt, 2006; Postle et al., 2007; Sosa Ponce et al., 2024) or with membrane-less nuclear foci such as nuclear speckles or nucleoli (Jacobsen et al., 2019; Morovicz et al., 2022; Vidalle et al., 2023). The lipids detected in the nucleus consist of sphingolipids, neutral lipids and glycerophospholipids, including polyphosphoinositides (PPIn) (Albi et al., 2003; Mate et al., 2006; Postle et al., 2007). PPIn (review on nomenclature (Michell et al., 2006)) are signalling lipids regulating a myriad of cellular processes not only from the cytoplasm but also from the nucleus (Jacobsen et al., 2019; Morovicz et al., 2022; Y. H. Wang & Sheetz, 2022; Vidalle et al., 2023). PPIn are phosphorylated derivatives of the glycerophospholipid phosphatidylinositol (PtdIns), which consists of two hydrophobic fatty acyl chains esterified to a glycerol backbone, itself conjugated via a phosphodiester linkage to a *myo*-inositol ring. The inositol ring can be reversely phosphorylated at the 3’, 4’ and 5’ hydroxyl groups, producing seven different PPIn, *i.e.* the monophosphorylated PtdIns3*P*, PtdIns4*P* and PtdIns5*P*, the bisphosphorylated PtdIns(3,4)*P*_2_, PtdIns(3,5)*P*_2_ and PtdIns(4,5)*P*_2_, and finally the triphosphorylated PtdIns(3,4,5)*P*_3_. Quantitatively, about 15% of all cellular PPIn are present in the nucleus (York & Majerus, 1994) and all, except PtdIns(3,5)*P*_2_, have been detected in this compartment via different methods (Cocco et al., 1987; Divecha et al., 1991; Mazzotti et al., 1995; Vann et al., 1997; Boronenkov et al., 1998; Gillooly et al., 2000; Clarke et al., 2001; Osborne et al., 2001; Watt et al., 2002; Jones et al., 2006; Lindsay et al., 2006; Kwon et al., 2010; Sarkes & Rameh, 2010; Yildirim et al., 2013; Kalasova et al., 2016; Karlsson et al., 2016). In contrast to cytoplasmic PPIn, which are embedded in membranes via their acyl chains, PPIn have been detected in membrane-less sites within the confines of the nucleus. How the hydrophobic groups are accommodated biophysically is however not entirely understood. Several studies have shown that the acyl chains could be shielded in the hydrophobic ligand pocket of the nuclear receptors, steroidogenic factor 1 and liver receptor homolog-1 (Krylova et al., 2005; Blind et al., 2012; Blind et al., 2014; Sablin et al., 2015), allowing the presentation of the hydrophilic inositol ring for its modification by PPIn metabolising enzymes (Blind et al., 2012). Alternatively, the acyl chains of PtdIns(4,5)*P*_2_ were suggested to be anchored on the myristoyl group conjugated to the transcription repressor BASP-1 (brain acid soluble protein 1) (Toska et al., 2012) but how these hydrocarbon chains are shielded is not clear. Lipid droplets have also been identified in the nucleus (Layerenza et al., 2013; Fujimoto, 2024) and may act as a platform where acyl chains could be buried through their monolayers.

A major pool of PtdIns(4,5)*P*_2_ is found in nuclear speckles, a nuclear sub-compartment involved in splicing and other mRNA metabolism (Galganski et al., 2017), together with the PtdIns(4,5)*P*_2_ metabolising enzymes, phosphatidylinositol-4-phosphate 5-kinase type Iα (PIP5K1A), phosphatidylinositol-5-phosphate 4-kinase type IIα/β (PIP4K2A/B) and phospholipase Cβ1 to a certain extent (Boronenkov et al., 1998; Ciruela et al., 2000; Osborne et al., 2001; Watt et al., 2002; Tabellini et al., 2003; Bultsma et al., 2010). A minor PtdIns(4,5)*P*_2_ pool is also found in nucleoli (Yildirim et al., 2013; Kalasova et al., 2016) or nuclear islets (Sobol et al., 2018). PtdIns(4,5)*P*_2_ was shown to regulate mRNA processing and splicing (Osborne et al., 2001; Mellman et al., 2008) and transcription via RNA polymerase I and II (Toska et al., 2012; Yildirim et al., 2013). PPIn are known to exert their function through their interaction with nuclear proteins via structured PPIn-binding domains (Di Lello et al., 2005) or lysine-rich polybasic regions (PBR) ((Gozani et al., 2003; Ahn et al., 2005; Huang et al., 2007; Viiri et al., 2009; Bidlingmaier et al., 2011; Lewis et al., 2011; Gelato et al., 2014; Stijf-Bultsma et al., 2015; Mazloumi Gavgani et al., 2021) and reviewed in (Jacobsen et al., 2019)). To gain further insight into the nuclear functions of PPIn, we have previously established quantitative interactomics methods to enrich for nuclear proteins representing potential PPIn effector proteins (Lewis et al., 2011; Mazloumi Gavgani et al., 2021). In combination with PtdIns(4,5)*P*_2_ pull-down, this approach led to the identification of DNA topoisomerase 2-alpha (TOP2A) as one of the PPIn effector protein (Lewis et al., 2011).

DNA topoisomerases consist of a family of enzymes that resolve DNA topological features including catenation and supercoiling occurring during replication, transcription and sister chromatid entanglement during mitosis, that would otherwise be deleterious (Pommier et al., 2022). TOP2A is one of the two mammalian type II isoforms with TOP2B, which catalyse double-strand breaks in one DNA helix, allowing the passage of the other DNA helix through the newly created gap before its ligation (Vos et al., 2011). Both TOP2A and TOP2B consist of an N-terminal ATPase domain, a central domain with catalytic activity and a C-terminal domain (CTD) with regulatory functions (Gardiner et al., 1998; McClendon & Osheroff, 2007). Both isozymes have 72 % identity but have the highest sequence divergence in their CTD and in the N-terminus (Jenkins et al., 1992). The CTD is not required for *in vitro* activity as truncated TOP2A and TOP2B are both still able to decatenate DNA, but is needed *in vivo* to support cellular growth (Meczes et al., 2008). Engineering chimeric enzymes with switched CTDs demonstrated differences between the two isozymes with regards to proliferation, activity, chromosome binding and DNA geometry recognition (Linka et al., 2007; McClendon et al., 2008; Meczes et al., 2008 ref update?). In particular, TOP2A prefers to relax positively supercoiled DNA, which accumulates in front of replication forks and this preference is believed to be due to positively charged amino acids located in the CTD which help recognise the different DNA supercoils (McClendon et al., 2005; McClendon et al., 2008).

TOP2A is ubiquitously expressed and plays important roles in DNA replication, transcription and chromatid segregation. The level of TOP2A expression is cell-cycle dependent (Drake et al., 1989; Woessner et al., 1991; Goswami et al., 1996), with lowest levels of expression in G1 and G0 phases and highest levels at late S, G2 and M phases of the cell cycle. During mitosis, TOP2A is associated with chromosomes (Meyer et al., 1997; Christensen et al., 2002), while in interphase, TOP2A resides within the nucleoplasm where it is associated with different foci, such as nuclear speckles (particularly when transcription is inhibited), the nucleolus and centromeric heterochromatin (Rattner et al., 1996; Meyer et al., 1997; Christensen et al., 2002; Agostinho et al., 2004). How TOP2A is targeted to different nuclear foci, is however, not understood. We previously identified an interaction between TOP2A and PtdIns(4,5)*P*_2_ or PtdIns5*P* (Lewis et al., 2011), and considering their common localisation in nuclear speckles, this led us to characterise their interaction and sub-cellular association in more detail. In this study, we have shown that TOP2A binds directly to PPIn with differing specificity and identified two polybasic regions (PBR) in its CTD, which contribute both to its interaction with PPIn and nucleolar localisation. Using immunolabeling, we have also shown that TOP2A co-localises with PtdIns(4,5)*P*_2_ in nuclear speckles upon RNA polymerase inhibition and to the chromatin upon the inhibition of PIP4K2A/B and subsequent reduction of PtdIns(4,5)*P*_2_.

## Materials and methods

### Reagents

The PIPK selective inhibitors for PI4P5K1A (ISA-2011B (#HY-16937) (Semenas et al., 2014)) and PI5P4K2 (PIP4K-IN-a131 (#HY-136310) (Kitagawa et al., 2017)) were purchased from MedChemExpress, dissolved in DMSO and stored at -80°C. α-amanitin (A2263) was from Sigma-Aldrich. Antibodies and the dilutions used in western blotting and immunostaining are listed in Table 1.

**Table 1.** List of antibodies used in this study. IMF: immunofluorescence staining, LOA: lipid overlay assay, RT room temperature and WB: Western immunoblotting. HRP: horseradish peroxidase. RRID: resource report ID

| Antibody target | Catalogue Number | Company name | RRID | Dilution |
| --- | --- | --- | --- | --- |
| DNA Topoisomerase 2α (clone Ki-S1) | MAB4197 | Merck Millipore | AB_2205862 | **IMF:** 1:500 |
| DNA Topoisomerase 2α [EP1102Y] | 52934 | Abcam | AB_883143 | **WB:** 1:2000 |
| Flag | F7425 | Sigma-Aldrich | AB_439687 | **IMF:** 1:160 |
| GAPDH (6C5) | sc-32233 | Santa Cruz Biotechnology | AB_627679 | **WB:** 1:5000 |
| GST-HRP | ab3416 | Abcam | AB_303783 | **LOA:** 1:30,000 |
| Histone H3 | 07-690 | Millipore | AB_417398 | **WB:** 1: 10,000 |
| Lamin A/C (E-1) | sc-376248 | Santa Cruz Biotechnology | AB_10991536 | **WB:** 1:10,000 |
| Nucleolin (D4C7O) | 14574 | Cell Signaling Technology | AB_2798519 | **IMF:** 1:1000 (o/n) or 1:100 1 h RT |
| Nucleophosmin (FC-61991) | 32-5200 | Thermo Fisher Scientific (Invitrogen) | AB_86912 | **IMF:** 1:500 |
| PARP1 (46D11) | 9542 | Cell Signaling Technology | AB_2160739 | **WB:** 1:2000 |
| PIP4K2A | 5527 | Cell Signaling Technology | AB_2722636 | **WB:** 1:1000 |
| PIP4K2B | 9694 | Cell Signaling Technology | AB_2164572 | **WB:** 1:1000 |
| PIP5K1A | 9693 | Cell Signaling Technology | AB_2164698 | **WB:** 1:1000 |
| PtdIns(4,5)$P_2$ (mouse IgM) clone 2C11 | sc-53412 | Santa Cruz Biotechnology | AB_630097 | **IMF:** 1:50 |
| α-tubulin | T5168 | Sigma-Aldrich | AB_477579 | **WB:**1: 20,000 |
| goat anti-mouse (IgM) Alexa594 | A-21044 | Thermo Fisher Scientific | AB_2535713 | **IMF:** 1:200 |
| goat anti-mouse (IgG) Alexa594 | A-11005 | Thermo Fisher Scientific | AB_2534073 | **IMF:** 1:200 |
| goat anti-rabbit (IgG) Alexa488 | A-11008 | Thermo Fisher Scientific | AB_143165 | **IMF:** 1:200 |
| anti-mouse (IgG2a) Alexa488 | A-21131 | Thermo Fisher Scientific | AB_2535771 | **IMF:** 1:200 |
| goat anti-mouse (IgG)-HRP | G-21040 | Thermo Fisher Scientific | AB_2536527 | **WB:**1: 10,000 |
| goat anti-rabbit (IgG)-HRP | G-21234 | Thermo Fisher Scientific | AB_2536530 | **WB:**1: 10,000 |

### Plasmids

The pGEX-6P-1-hTOP2A-CTD (containing amino acids 1171-1531) construct was provided by Susan P.C. Cole (Queen’s University, Canada). hTOP2A-Δ1365-1431-CTD and TOP2A-CTD PBR deletion mutants (ΔM1-ΔM6) were generated by overlap extension PCR using pGEX-6P-1-hTOP2A-CTD as template and cloned in-frame at the 3’ end of Glutathione-S-Transferase (GST) using the XmaI (ΔM3-4 and Δ1365-1431) or SmaI (ΔM1, ΔM2, ΔM5 and ΔM6) and XhoI sites of pGEX-4T1. hTOP2A-CTD and hTOP2A-CTD-ΔM3-4 were sub-cloned into pEGFP-C2 via XhoI and XmaI 3’ to GFP. GFP tagged pEGFP-C2-hTOP2A-CTD-ΔM2 and pEGFP-C2-hTOP2A-CTD-ΔM2 + M3-4 were generated from pEGFP-C2-hTOP2A-CTD and pEGFP-C2-hTOP2A-CTD-ΔM3-4 respectively by deletion site directed mutagenesis. FLAG-tagged full-length TOP2A WT (Flag-FL-TOP2A-WT) was prepared by PCR cloning from pCMV-IRES-GFP-TOP2A (obtained from W. Beck) into the NotI and SmaI restriction sites of pFLAG-CMV-2 vector (Sigma-Aldrich, USA). Flag-FL-TOP2A-ΔM3-4 was generated by overlap extension PCR using Flag-FL-TOP2A-WT as template. Flag-FL-TOP2A-ΔM2 and Δμ2+M3-4 were generated by deletion site directed mutagenesis using Flag-FL-TOP2A-WT as template. In all cases, mutations were confirmed by sequencing.

### Purification of GST fusion proteins

BL21(DE3)-RIL *Escherichia coli* cells were transformed with the pGEX-6P-1-TOP2A-CTD or pGEX-4T1-TOP2A-CTD deletion constructs and bacterial cultures were induced with 100 µM IPTG overnight at 25°C or at 37°C for 3 h. The next day, cells were harvested by centrifugation and the pellet was resuspended in PBS containing 1% Triton, 10 mM DTT and a protease inhibitor cocktail (1:100; Sigma cat P8849). The cell lysates were snap frozen at - 80°C and thawed the next day at 37°C. Sonication was carried out for 3 x 30 sec followed by centrifugation, and the supernatant was batch-bound to Glutathione Sepharose 4B for 1 h with end-on-end rotation at 4°C. GST-TOP2A-CTD fusion proteins were eluted with 50 mM Tris, pH 8, containing 300 mM NaCl, 10 mM reduced glutathione and 10 mM DTT. The eluted proteins were dialysed against PBS for 1 h at 4°C. After dialysis, the purified proteins were aliquoted and stored at -80°C.

### Protein-lipid overlay assay

Lipid overlay assays were performed as described previously (Karlsson et al., 2016) using PPIn Strips− (Echelon Biosciences Inc.) and 0.5 µg/ml of GST-fused recombinant proteins in 5 mL of blocking buffer (PBS, pH 7.4, 3% fatty acid-free BSA). Each protein-lipid overlay assay was done at least thrice. The signals from lipid blots were quantified using Image J, v1.52p (http://imagej.net/ij RRID:SCR_003070).

### Cell culture and transfection

HeLa cells (American Type Culture Collection, RRID:CVCL_0030) were cultured in DMEM-Hi glucose medium and TIG-1 normal human foetal lung fibroblast cells (Coriell Institute For Medical Research, Camden, NJ, USA, RRID:CVCL_0560) in Eagle’s Minimum Essential Medium (MEM) with Earle’s salts and non-essential amino acids. Both media were supplemented with antibiotics (Penicillin-Streptomycin) and 10% FBS and maintained at 37°C with an atmosphere of 5% CO_2_ (growth medium). For GFP-tagged TOP2A CTD experiments, HeLa cells were grown on glass coverslips in 6-well plates for a day before they were transfected with 2 µg pEGFP-C2-TOP2A-CTD constructs using 5 µL Lipofectamine 2000 transfection reagent or 1 µg Flag-FL-TOP2A constructs and 3 µL X-tremeGENE 9 or Lipofectamine 3000, according to manufacturer’s instructions. Complexes were incubated with cells in serum and antibiotic free medium, which was replaced 4 h later to growth medium for a further 20 h.

### Whole cell extract preparation and subcellular fractionation

HeLa cells were treated with PIPK inhibitors for 24 h and both attached and detached cells were collected, washed in PBS and pooled. Cell pellets were lysed in either radioimmunoprecipitation assay buffer (50 mM Tris-HCl pH8.0, 0.5% deoxycholic acid, 150 mM NaCl, 1% NP-40 and 0.1% SDS, supplemented with 5 mM NaF, 2 mM Na_3_VO_4_ and 1x Sigma Protease Inhibitor Cocktail) or in boiling 1% SDS buffer (1% SDS, 1 mM Na_3_VO_4_). Cell extracts obtained with 1% SDS lysis buffer were sonicated with 30% amplitude, 6x 5 sec ON-5 sec OFF). Nuclear fractionation was carried out according to (Karlsson et al., 2016) with some brief changes. Cells were grown in 10 cm plates up to 90% confluency and washed with ice cold PBS followed by a quick rinse with cold buffer A (10 mM HEPES pH 7.9, 1.5 mM MgCl_2_, 10 mM KCl). The cells were then incubated with buffer A containing 1% Igepal, 0.5 mM DTT, 2 mM Na_3_VO_4_ and Protease Inhibitor cocktail for 10 min on ice and centrifuged at 600 g for 5 min at 4°C. The supernatants were collected as the cytoplasmic fractions. The nuclear pellets were quickly rinsed twice with buffer A and centrifuged at 600 g for 6 min at 4°C each time. The final nuclear pellets were lysed with 200 μL RIPA lysis buffer and vortexed. After 10 min of incubation, the nuclear extract was centrifuged at 13 000 g for 5 min at 4°C and the supernatant collected. To isolate the chromatin enriched fractions, the nuclear pellets were resuspended with buffer B (3 mM EDTA, 0.2 mM EGTA, 1 mM DTT, 1:100 protease inhibitor cocktail), incubated for 30 min on ice (Mendez & Stillman, 2000) and centrifuged for 5 min, 1700 g at 4°C. The resulting pellets, corresponding to insoluble chromatin were washed quickly in Buffer B and finally resuspended in Laemmli buffer (without bromophenol blue) and sonicated 2x 15 sec.

### Western immunoblotting

Total protein concentrations were measured with the Pierce™ BCA Protein Assay Kit. Protein samples were boiled for 5 min in the presence of Laemmli sample buffer, equal amount of proteins were resolved on 10 % SDS-PAGE gel and transferred onto 0.45 µm thick nitrocellulose membranes. The membranes were blocked with 5 % skimmed milk in TBS-T (50 mM Tris pH 7.5, 150 mM, NaCl, 0.05% Tween 20) and incubated with primary antibodies overnight at 4°C or for 1 h at room temperature (see Table 1). This was followed by incubation of the membranes with horseradish peroxidase-conjugated secondary antibodies at 1:10,000 dilution for 1 h, at room temperature. The membranes were washed 6 x 5 min with PBS-T following each antibody incubation. The immunoreactive bands were visualised by enhanced chemiluminescence using SuperSignal West Pico Chemiluminescent Substrate, detected with a BioRad ChemiDocTM Xrs+ and quantified using image j, v1.52p (http://imagej.net/ij). (Schindelin et al., 2012).

### Immunofluorescence staining and microscopy

Cells were grown on glass coverslips in 6-well plates and were fixed the day after in 3.7% paraformaldehyde in PBS, pH 7.4, for 10 min at room temperature. Subsequently, cells were permeabilized with 0.25% Triton in PBS for 10 min, which allows to detect detergent-resistant PPIn in the nucleus (Osborne et al., 2001; Mortier et al., 2005). Cells were blocked with 3% BSA, 0.05% Triton-X100 in PBS for 1 h at room temperature. The cells were then incubated with primary antibodies (anti-PtdIns(4,5)*P*_2_) overnight at 4°C followed by goat anti-mouse (IgM) Alexa594 for 1 h at room temperature. Cells were then incubated with primary anti-TOP2A antibody (either Abcam ab52934 or Merck Millipore KiS1 (Boege et al., 1995)), for 1 h at room temperature followed by incubation with secondary antibodies goat anti-rabbit (IgG) Alexa488 or goat anti-mouse (IgG2a) Alexa488 respectively for 1 h at room temperature. For the RNA Polymerase II inhibition experiments, cells were treated with solvent (DMSO) or 50 μg/ml α-amanitin for 5 h at 37°C before proceeding with immunolabelling as described above. GFP-tagged TOP2A-CTD transfected cells were processed for immunostaining as above and incubated with anti-nucleolin for 1 h at room temperature, followed by incubation with secondary antibody goat anti-rabbit (IgG)-Alexa594 for 1 h at room temperature. Flag-tagged FL-TOP2A transfected cells were stained with anti-Flag antibody overnight and goat anti-rabbit IgG-Alexa488 followed by anti-nucleophosmin and goat anti-mouse IgG-Alexa 594. Cells were washed 3-4 times in PBS-Tween-20 (0.05%) after each antibody incubation and mounted in ProLong Gold antifade reagent with 4′,6-diamidino-2-phenylindole (DAPI). Fluorescent images were captured using either a Leica TCS SP2 confocal microscope or a Leica DMI6000B fluorescent microscope with x40 or x100 objectives. Images were processed with the Leica application suite X v3.7.4 and immunofluorescence intensity of PtdIns(4,5)*P*_2_ was measured in the nucleus using Fiji (https://imagej.net/software/fiji/, RRID:SCR_002285) (Schindelin et al., 2012)).

### Statistical analyses

Three to five biological repeats were performed for each experiment and GraphPad Prism (RRID:SCR_002798) version 10.0.2 was used for making graphs and performing statistical analyses. The exact statistical tests performed are indicated in the figure legends.

## Results

### TOP2A interacts directly with polyphosphoinositides and colocalises with PtdIns(4,5)*P*2 in nuclear speckles in response to RNA polymerase II inhibition

In a previous study, we identified TOP2A as a potential binding partner for nuclear PPIn and showed that the CTD of TOP2A could bind to PtdIns(4,5)*P*_2_ as well as PtdIns5*P*, and less so to PtdIns3*P* (Lewis et al., 2011). To determine the PPIn-binding specificity of TOP2A-CTD, we expressed this domain as a GST-fusion protein and tested its interaction with a wider range of phospholipids in lipid overlay assays (Figure 1A). GST-TOP2A-CTD was found to bind to all PPIn and phosphatidic acid (PA), but not to any of the other phospholipids tested, while lipid binding was not detected for GST alone (Figure 1A). Significant interactions were observed for PtdIns4*P*, PtdIns(3,4)*P_2_*, PtdIns(3,5)*P*_2_, PtdIns(4,5)*P*_2_, and PtdIns(3,4,5)*P*_3_, while those with PtdIns3*P* and PtdIns5*P* and PA were more variable (Figure 1B).

**Figure 1:**
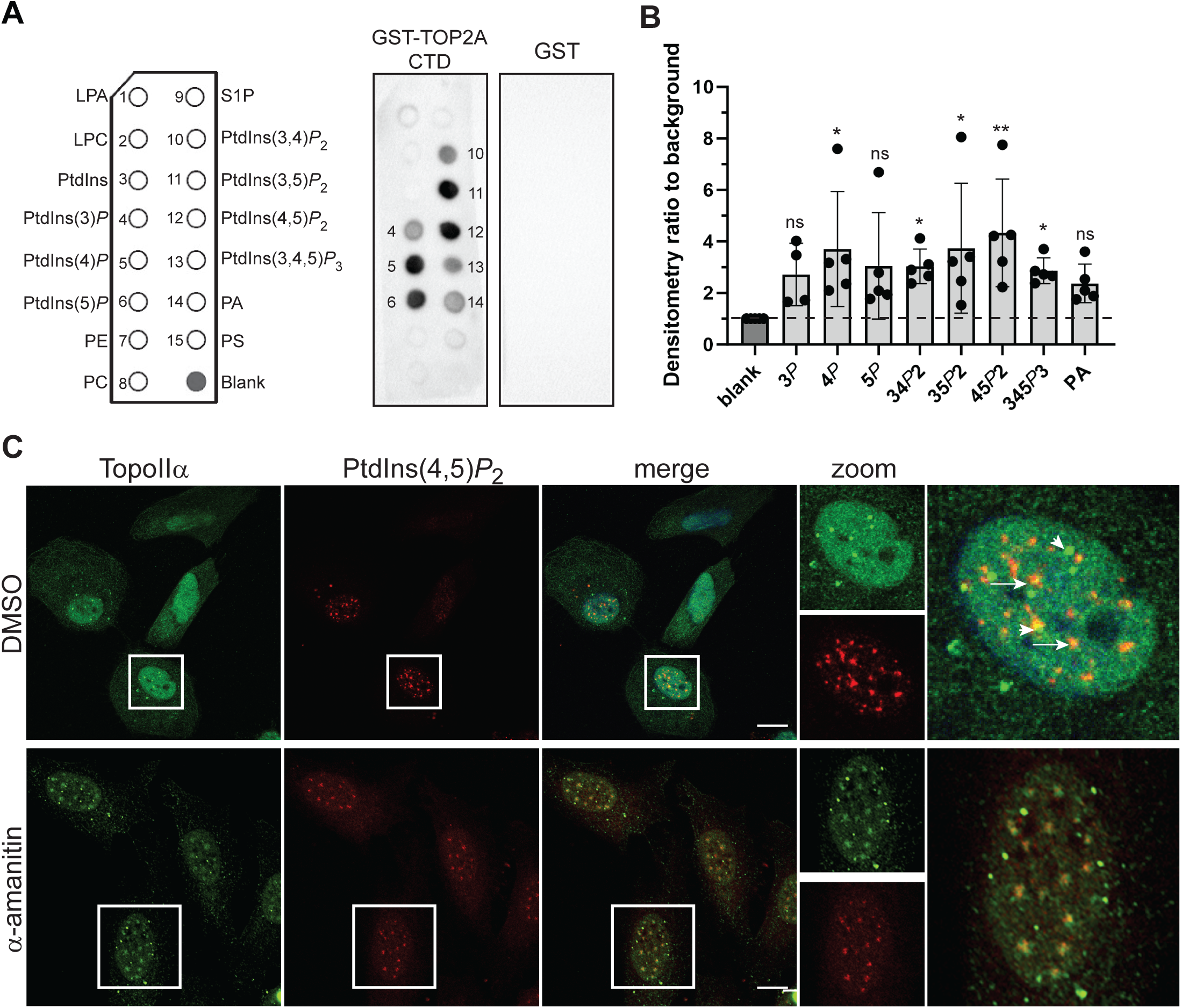
TOP2A binds to PPIn and associates with PtdIns(4,5)*P*2 in the nucleus. ***A, Left***: PIP strip schematic (www.echelon-inc.com) showing the positions of the following glycerophospholipids: lysophosphatidic acid (LPA), lysophosphocholine (LPC), phosphatidylinositol (PtdIns), PtdIns3*P*, PtdIns4*P*, PtdIns5*P*, phosphatidylethanolamine (PE), phosphatidylcholine (PC), sphingosine-1-phosphate (S1P), PtdIns(3,4)*P*2, PtdIns(3,5)*P*2, PtdIns(4,5)*P*2, PtdIns(3,4,5)*P*3, phosphatidic acid (PA), phosphatidylserine (PS) and blank. ***Right***, 0.5 µg/ml of GST-TOP2A-CTD or GST were incubated with PPIn strips and protein-lipid interactions were detected with anti-GST-HRP and chemiluminescence. Lipids bound are labelled accordingly to the schematic on the left. ***B***, Quantification of the protein-lipid interaction signals measured from 5 experiments shown as means +/- SDs of the densitometry ratios related to background signal. ns, not significant, * *P*<0.05, ** *P*<0.01 (One-way ANOVA, non parametric, Kruskal-Wallis multiple comparison test). ***C***, Asynchronous and α-amanitin (50 µg/mL, 5 h) treated HeLa cells stained with anti-TOP2A KiS1 mAb and anti-PtdIns(4,5)*P*2 IgM antibody; white squares indicate nuclei that are enlarged in the zoom panel. Scale bar, 10 µm. Representative of three independent experiments.

TOP2A has previously been described to accumulate in nuclear speckles upon the inhibition of transcription (Agostinho et al., 2004), a site where PtdIns(4,5)*P*_2_ was also shown to accumulate in several independent studies (Boronenkov et al., 1998; Osborne et al., 2001; Watt et al., 2002). We therefore tested whether TOP2A colocalised with PtdIns(4,5)*P*_2_ in these structures by immunofluorescence studies in HeLa cells in the absence or presence of RNA polymerase II inhibitor (Figure 1C). In untreated cells, TOP2A exhibited a punctate nucleoplasmic staining pattern with a few intense foci (Figure 1C, arrow heads), which were described to contain TOP2A and the centromere protein CENP (Agostinho et al., 2004). In contrast, PtdIns(4,5)*P*_2_ stained as strong irregular foci, called nuclear speckles (Figure 1C), when using the same permeabilization method to detect detergent-resistant PPIn in the nucleus as in previous studies (Boronenkov et al., 1998; Osborne et al., 2001; Watt et al., 2002). In some cells, a partial overlap could be observed between PtdIns(4,5)*P*_2_ and TOP2A puncta or other smaller and less intense foci (Figure 1C, full arrows), particularly in cells with the most intense PtdIns(4,5)*P*_2_ signal. RNA polymerase II inhibition with α-amanitin resulted in the reorganisation of PtdIns(4,5)*P*_2_-positive speckles to fewer structures, as previously reported (Osborne et al., 2001). Concomitantly, TOP2A accumulated in PtdIns(4,5)*P*_2_-positive speckles in most cells (Figure 1C). These observations were confirmed in another cell line, a normal human fibroblast cell line, TIG-1 (Supplementary Figure S1). Taken together, recombinant TOP2A-CTD was shown to interact directly with several PPIn, including PtdIns(4,5)*P*_2_, while endogenous TOP2A colocalises with speckle-associated PtdIns(4,5)*P*_2_ upon RNA polymerase II inhibition.

### TOP2A interacts with polyphosphoinositides via two polybasic regions located in the C-terminal domain

To identify the PPIn interaction site, we analysed the CTD sequence and noticed that TOP2A does not harbour PPIn-binding modules such as PH, PX or FYVE domains, but contains six lysine-rich polybasic regions (PBR), following the K/R motif sequence K/R-(X_n=3-7_)-K-X-K/R-K/R (Martin, 1998) (Figure 2A). The lysine residues within these PBR are particularly well conserved in vertebrates ((Endsley et al., 2024), supplementary Figure S2), suggesting important function. To determine the contribution of each PBR in mediating PPIn-binding, we generated deletion mutants lacking each PBR, named motifs (M): M1 (1196-KGGKAKGKK-1204), M2 (1228-KAEAEKKNKK-1237), M3-4 (1259-KQRLEKKQKREPGTKTKK-1276), M5 (1283-KPIKKGKK-1290) or M6 (1454-KPDPAKTKNRR-1464). M3 and 4 overlapped and were hence deleted together (Figure 2A). The deletion mutants were expressed as GST-fusion proteins and tested for their lipid binding properties in lipid overlay assays, compared to WT and a deletion mutant lacking amino acid residues from a region located between M5 and M6 (Δ1365-1431) (Figure 2B). The loss of M2 or M3-4 consistently almost abolished TOP2A-CTD binding to PPIn (Figure 2B). Deletion of M6 also led to a decrease in PPIn binding, albeit to a lesser extent compared to M2 or M3-4, while deletion of M1, M5 or the region spanning amino acids 1365-1431 did not affect TOP2A-CTD binding to PPIn. Taken together, these results indicate that the M2 and/or M3-4 of TOP2A-CTD are necessary to bind PPIn and their importance is reflected by their conservation in vertebrates (Figure 2C).

**Figure 2:**
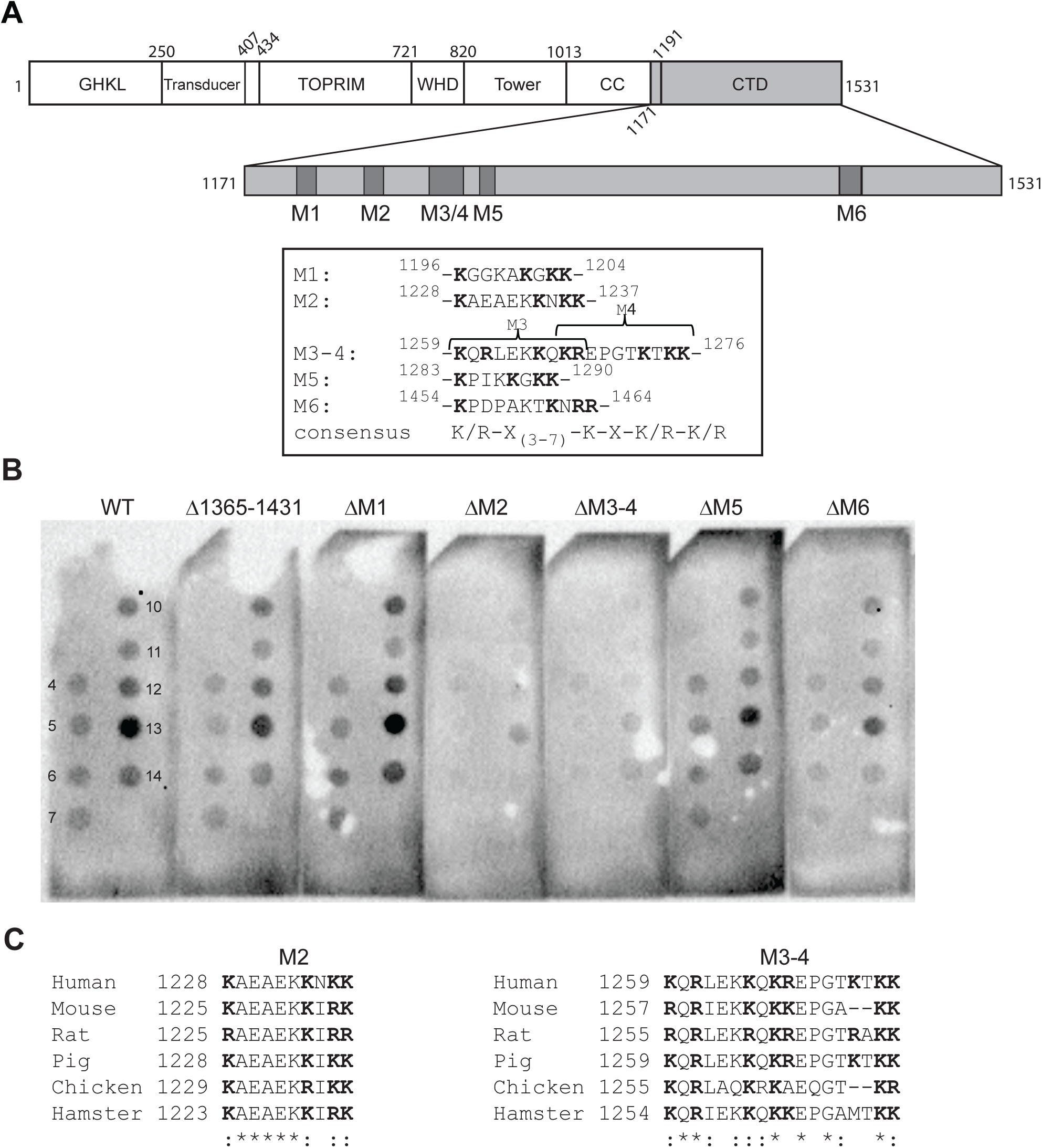
TOP2A binds to PPIn via two polybasic regions in its C-terminal domain. ***A***, Schematic of TOP2A domain structure according to Broeck *et al* (Vanden Broeck et al., 2021). The positions of the polybasic regions (PBR) motifs M1-M6 are shown as black bars in the CTD and their respective sequences are shown in the box below. Basic residues are highlighted in *bold*. ***B***, 0.5 µg/ml of the indicated GST-TOP2A-CTD fusion proteins: WT, Δ1365-1431, ΔM1, ΔM2, ΔM3-4, ΔM5, and ΔM6 were incubated with PPIn strips and interactions were detected with anti-GST-HRP and chemiluminescence. Sequence deleted in Δ1365-1431: SNKELKPQKSVVSDLEADDVKGSVPLSSSPPATHFPDETEITNPVPKKNVTVK KTAAKSQSSTSTTG. Representative of four independent experiments. ***C***, Sequence alignment of M2 and M3-4 of human TOP2A (P11388-1) compared to other vertebrate species, mouse (Mus musculus, Q01320-1), rat (Rattus norvegicus, P41516-1), pig (Sus scrofa, O46374-1), chicken (Gallus gallus, O42130-1) and Chinese hamster (Cricetulus griseus (Cricetulus barabensis griseus), P41515-1) , taken from the whole alignment results of the CTD in Supplemental Figure S2.

### The polybasic regions M2 and M3-4 contribute to the nucleolar localisation of TOP2A

PBR found in nuclear PPIn effector proteins have previously been shown to exhibit dual functions by acting as 1) PPIn interaction sites and 2) nuclear or nucleolar localisation signals (Viiri et al., 2009; Toska et al., 2012; Karlsson et al., 2016). Analysis of the PBR M2 and M3-4 using the nucleolar sequence detector (NoD, (Scott et al., 2011)), revealed that they harbour putative nucleolar localisation signals (Figure 3A). In addition, M2 is located within a previously reported weak bipartite NLS, spanning amino acids 1259-1296 (Mirski et al., 1999). Since the deletion of the M2 and M3-4 PBR motifs resulted in the loss of PPIn binding, we tested the effect of their deletion on the localisation of TOP2A. To this end, we transiently transfected HeLa cells with EGFP-TOP2A-CTD WT, the single deletion mutants ΔM2 and ΔM3-4, as well as the ΔM2-ΔM3-4 double mutant and immunostained them with the nucleolar marker nucleolin (Figure 3B). TOP2A-CTD WT localised in the nucleoplasm and a strong signal was observed in the nucleolus in 92 % of EGFP-positive cells (Figure 3B-C). The remaining cells showed a diffuse nuclear pattern. Single deletion of either of M2 or M3-4 led to a significant decrease in the percentage of cells with nucleolar CTD and a concomitant increase of the diffuse pattern. The M2 mutant did not lead to cytoplasmic localisation of the CTD. The ΔM3-4 mutant also exhibited a ring pattern around nucleoli, which colocalised with nucleolin. When both PBR were deleted, a further increase of both the diffuse and ring patterns with a concomitant decrease of the nucleolar localisation of TOP2A-CTD was observed. EGFP alone showed signal both in the cytoplasm and nucleus (data not shown). To check the effect of each motif on the localisation of the full length protein, Flag-tagged constructs for the WT and the three deletion mutants were generated. Following transient transfection, HeLa cells were stained to detect the nucleolar protein nucleophosmin (Figure 3D). Overexpression of the full length TOP2A WT resulted in a higher proportion of cells showing a diffuse signal in the nucleus and a lower with nucleolar localisation than was observed for the isolated CTD (Figure 3D-E). Deletion of either M2 or M3-4 resulted in a significant decrease in the nucleolar localisation and a concomitant increase in the proportion of cells devoid of nucleolar signal (Figure 3E), similarly to the CTD alone (Figure 3C). Too few cells survived when both PBR were deleted in the full length protein (data not shown). The full length TOP2A did not tend to display the ring pattern compared to the CTD only. In conclusion, the M2 and M3-4 PBR contribute to the nucleolar localisation of TOP2A.

**Figure 3:**
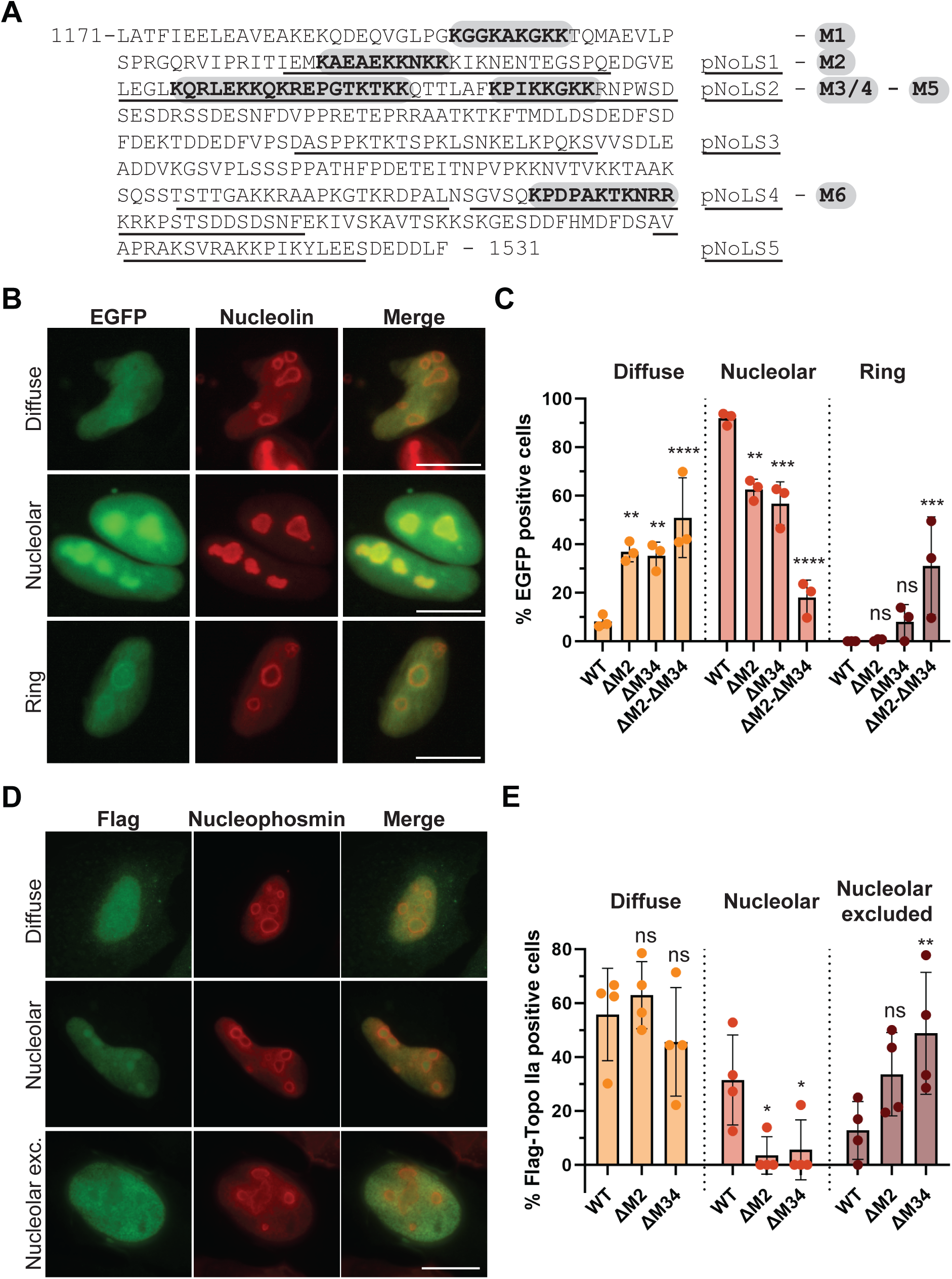
Loss of PBR motifs M2 and M3-4 prevents TOP2A nucleolar localization. ***A***, Amino acid sequence of the C-terminal domain (CTD) of human TOP2A, showing the polybasic motifs (grey) and the putative nucleolar localisation sequence (pNoLS, underlined). ***B***, HeLa cells were transfected with EGFP-CTD-TOP2A WT, ΔM2, ΔM3-4, ΔM2/ΔM3-4 or empty pEGFP-C2 vector, stained with anti-nucleolin antibody and imaged by epifluorescence microscopy. EGFP positive cells were classified according to the three following localisation patterns: *Nucleolar* – accumulation of EGFP signal in nucleoli compared to the surrounding nucleoplasm; *Diffuse* – no marked accumulation of EGFP signal in nucleoli and *Ring* – no accumulation of EGFP signal in nucleoli compared to nucleoplasm, but a ring around the nucleolus colocalising with nucleolin is visible, indicating the granular component of the nucleolus. EGFP alone was also present in the cytoplasm in addition to the nucleus (not shown). Example images for each localisation pattern, scale bar 10 μm. ***C***, Quantification of patterns from *B* from 3 independent experiments. Graphs of mean percentages ± SDs. ns: not significant, \*\**P*<0.01, \*\*\**P*<0.001, \*\*\*\**P*<0.0001 compared to WT (two-way ANOVA, Dunnett’s post test). ***D***, HeLa cells were transfected with Flag-FL-TOP2A WT, ΔM2, ΔM3-4, stained with anti-Flag and anti-nucleophosmin antibodies and imaged by epifluorescence microscopy. Flag positive cells were classified according to the following localisation patterns: Nucleolar and Diffuse as in *B*, as well as Nucleolar excluded -nucleoli devoid of Flag signal. Example images for each localisation pattern, scale bar 10 μm. ***E***, Quantification of patterns from *C* from 4 independent experiments. Graphs of mean percentages ± SDs, ns: not significant, \**P*<0.05, \*\**P*<0.01 compared to WT (two-way ANOVA, Dunnett’s test).

### Inhibition of phosphatidylinositol-5-phosphate 4-kinase type IIα/β enhances the association between TOP2A and the chromatin

PtdIns(4,5)*P*_2_ interaction has been shown to regulate the protein levels and stability of mutant p53 (Choi et al., 2019) and of the nuclear factor erythroid 2-related factor 2 (Chen et al., 2023). Considering the interaction of TOP2A with PPIn, and notably PtdIns(4,5)*P*_2_, we aimed to manipulate the levels of PtdIns(4,5)*P*_2_ by using selective inhibitors targeting nuclear PIPK enzymes and test the effect on TOP2A protein levels. We first verified the nuclear presence of PIPK enzymes in HeLa cells, which generate PtdIns(4,5)*P*_2_ from PtdIns4*P* (PIP5K1A) and from PtdIns5*P* (PIP4K2A and B) (Figure 4A-B), and which were previously reported to be present in the nucleus in other cell types (Boronenkov et al., 1998; Ciruela et al., 2000; Mellman et al., 2008; Bultsma et al., 2010). Using cell fractionation and western immunoblotting, all three enzymes were shown to be present in both the cytoplasm and nuclear fractions (Figure 4A). We next tested the effect of selective inhibitors of type I and II PIPK enzymes by measuring PtdIns(4,5)*P*_2_ nuclear signal intensities in HeLa cells using immunofluorescence (Figure 4C-D). Inhibition of PI5P4K2 with PIP4K-IN-a131, but not of PI4P5K1A with the selective inhibitor ISA-2011B, led to a significant decrease in nuclear PtdIns(4,5)*P*_2_ signal mean intensity (Figure 4C-D). Treatment with either inhibitor led to the detachment of a large proportion of cells, and induced cell death as shown by PARP1 cleavage (Figure 4E), which was consistent with previous studies in other cancer cell types (Kitagawa et al., 2017; Sarwar et al., 2019). The protein levels of TOP2A were next examined following the same treatments in total cell extracts (Figure 4F). Control cells exhibited low protein levels and the inhibition of PI4P5K1A did not show any significant change (Figure 4F). In contrast, inhibition of PI5P4K2A/B led to a dose-dependent and significant increase in TOP2A levels (Figure 4F-G). To try to understand the difference in TOP2A protein levels under these conditions, we examined how TOP2A was distributed in the nucleoplasm and the chromatin enriched fraction (CEF). A high level of TOP2A was detected in the nucleoplasm under all conditions, while a significant increase in the association of TOP2A with the chromatin was observed upon the inhibition of PI5P4K2A/B, but not of PI4P5K1A (Figure 4H-I). These results suggest that TOP2A is strongly associated with the chromatin upon the inhibition of PIP4K2A/B and may indicate an increase in TOP2A-DNA cleavage complexes. This would imply that different pools of TOP2A associate with the nucleoplasm and the chromatin in response to changes in the activity of PI5P4K2A/B.

**Figure 4:**
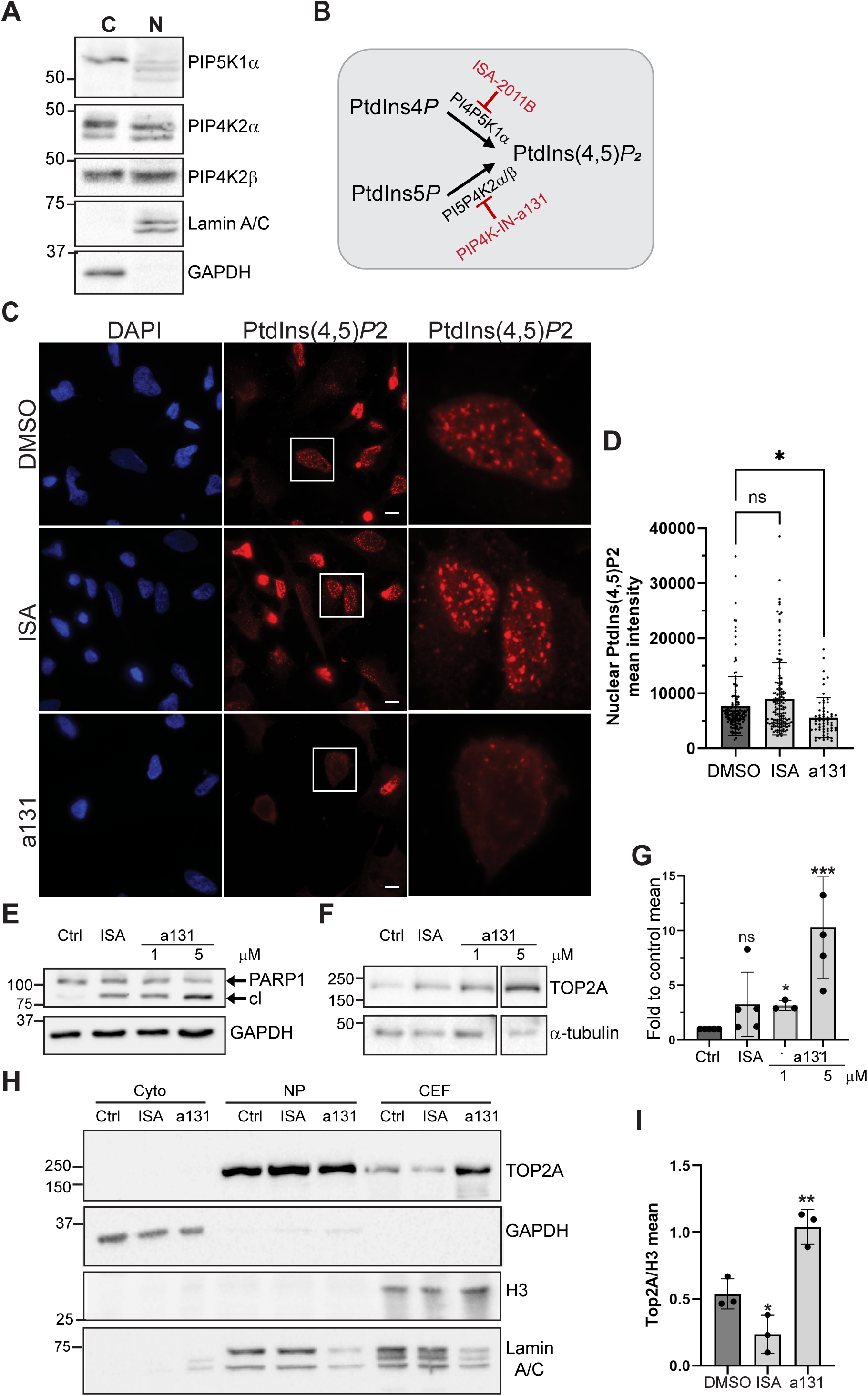
Inhibition of phosphatidylinositol-5-phosphate 4-kinase type IIα/β increase TOP2A association with the chromatin. ***A***, Asynchronous HeLa cells were fractionated, and cytoplasmic and nuclear extracts were resolved by SDS-PAGE and Western blotted using the indicated antibodies. GAPDH and Lamin A/C were used as fraction purity control for the cytoplasm and nucleus respectively. Representative blots of 3 independent experiments. ***B***, Schematic of the metabolism of phosphatidylinositol(4,5)bisphosphate (PtdIns(4,5)*P*2) in the nucleus generated from PtdIns4*P* via phosphatidylinositol 4-phosphate 5-kinase type-1 alpha (PI4P5K1α) and from PtdIns5*P* via phosphatidylinositol 5-phosphate 4-kinase type-2 alpha and beta (PI5P4K2α/β), as well as PIPK type I and II inhibitors. ***C-D***, Asynchronous HeLa cells were treated with 20 µM ISA-2011B (ISA) or 5 µM PIP4K-IN-a131 (a131) for 24 h and immunostained with anti-PtdIns(4,5)*P*2 antibodies and labelled with DAPI. Scale bar 10 µm. ***D***, Quantification of nuclear PtdIns(4,5)*P*2 intensity, n=4, ns, not significant, \**P*<0.05, (one-way ANOVA, Dunnett’s post test). ***E-F***, Asynchronous HeLa cells were treated with 20 µM ISA or 1-5 µM a131 for 24 h. Detached and attached cells were pooled, 20 µg protein was resolved by SDS-PAGE and blotted with the indicated antibodies. cl: cleaved PARP1. The separate blot boxes shown in *F* are from the same membrane and SDS-PAGE. ***G***, Quantification of TOP2A/α-tubulin ratio mean fold to Ctrl (DMSO), n=3-4, \**P*<0.05, \*\*\**P*<0.001, (one-way non-parametric ANOVA, Kruskal-Wallis test uncorrected Dunn’s test). ***H***, Asynchronous HeLa cells were treated with 20 µM ISA or 5 µM a131 for 24 h and subjected to subcellular fractionation to isolate soluble (nucleoplasm -NP) and insoluble chromatin fractions (chromatin enriched fraction – CEF). 20 µg from each fraction were resolved by SDS-PAGE and blotted with the indicated antibodies. ***I***, Quantification of TOP2A/histone H3 ratio from the CEF. n=3, \**P*<0.05, \*\**P*<0.01, (one-way ANOVA, uncorrected Fisher’s LSD).

## Discussion

We have previously identified TOP2A as a PPIn interacting protein and binding was demonstrated with PtdIns(4,5)*P*_2_ and PtdIns5*P in vitro* (Lewis et al., 2011). In this study, we mapped the interaction sites and identified two lysine-rich PBR located in the proximal end of the CTD, implying an electrostatic-mode of interaction with the phosphate groups of the inositol headgroup. This mode of interaction is consistent with previous studies that our group and others have demonstrated for several other nuclear proteins ((Skare & Karlsson, 2002; Gozani et al., 2003; Ahn et al., 2005; Kaadige & Ayer, 2006; Okada et al., 2008; Viiri et al., 2009; Gelato et al., 2014; Stijf-Bultsma et al., 2015; Karlsson et al., 2016; Mazloumi Gavgani et al., 2021), and reviewed in (Jacobsen et al., 2019)). The two PPIn-binding PBR identified in TOP2A, M2 and M3, are well conserved in vertebrates (Figure 2C) in contrast to the rest of the CTD which has low conservation ((Vanden Broeck et al., 2021), supplementary Figure S2), suggesting important functions. The analysis of the multiple alignment of 105 species demonstrated a high frequency of occurrence of lysine residues located in these particular regions ((Endsley et al., 2024), see also supplementary Figure S2), further highlighting the functional importance of these charged clusters. Structurally, how these motifs are presented to PPIn is still unclear since most of the CTD is predicted to be disordered and flexible (Vanden Broeck et al., 2021). None of the reported crystal structures include the CTD (Wendorff et al., 2012; Wang et al., 2017) and the recent cryo-EM study of the full length TOP2A was not able to detect any density from aa 1191 onwards (Vanden Broeck et al., 2021). In terms of the potential function of this interaction, PPIn may affect the activity of TOP2A, considering that we previously showed a decrease in TOP2A decatenation activity *in vitro* by bisphosphorylated PPIn and PtdIns(3,4,5)*P*_3_ (Lewis et al., 2011). Interestingly, the basic residues clustered in the CTD of TOP2A have previously been shown to be important for the activity of both the human and rat enzyme (McClendon et al., 2008; Yasuda et al., 2021). In particular, TOP2A-RNA interaction was shown to inhibit TOP2A catalytic activity *in vitro* (Park et al., 2008). In a more recent study of the rat TOP2A, the sequence encompassing residues 1192-1289 (corresponding to residues 1195-1293 in human TOP2A, spanning all motifs except M6) was necessary for its interaction with cellular RNA and this interaction led to the inhibition of relaxation activity of TOP2A (Yasuda et al., 2021). PPIn may compete with RNA for the interaction with the PBR of this region and may therefore regulate TOP2A activity. Alternatively, considering the effect of PPIn on decreasing TOP2A activity (Lewis et al., 2011), PPIn and RNA may interact together with TOP2A and lead to the inhibition of TOP2A. PPIn-RNA interaction has indeed been demonstrated in a few studies. A long noncoding (Lnc) RNA LINK-A was shown to interact with PtdIns(3,4,5)*P*_3_ and this interaction enhanced the association of PtdIns(3,4,5)*P*_3_ with AKT (Lin et al., 2017). A recent study established a method to map RNA bound to PPIn and identified RNA transcripts, including LncRNAs pulled down by PtdIns(4,5)*P*_2_ by RNAseq, suggesting that RNA-PPIn interaction may be more widespread as a type of biomolecular interaction (Bayona-Hernandez et al., 2023; Miladinovic et al., 2024). Whether this applies to TOP2A remains to be investigated.

TOP2A was shown in this study to accumulate to PtdIns(4,5)*P*_2_-rich nuclear speckles upon the inhibition of transcription, which reconciled previous isolated observations, showing PtdIns(4,5)*P*_2_ (Boronenkov et al., 1998; Osborne et al., 2001; Tabellini et al., 2003; Elong Edimo et al., 2011) and TOP2A (Agostinho et al., 2004) in these foci. Nuclear speckles are DNA poor and RNA-rich and TOP2A may be recruited to these sites to store the enzyme in its inactive form, bound to RNA and/or PtdIns(4,5)*P*_2_. Lack of activity associated with TOP2A in nuclear speckles has indeed been inferred previously, due to the depletion of TOP2A in DNA-poor sites, such as nuclear speckles and nucleoli, upon the addition of TOP2 inhibitors that traps the enzyme at active sites of the chromatin (Christensen et al., 2002; Agostinho et al., 2004). Moreover, since TOP2A can be inhibited by PPIn, including PtdIns(4,5)*P*_2_ *in vitro* (Lewis et al., 2011), it is possible that the association of PtdIns(4,5)*P*_2_ with TOP2A could lead to the presence of the inactive enzyme in the nuclear speckles. This may be consistent with the results of this study showing the significant decrease of TOP2A in the nucleoplasmic fraction and the concomitant increase in the enriched chromatin fraction upon the inhibition of PI5P4K2, decreasing a PI5P4K2-specific pool of PtdIns(4,5)*P*_2_. In contrast, the inhibition of PI4P5K1 decreased the level of TOP2A in the CEF. These results, combined with previous studies (Christensen et al., 2002; Agostinho et al., 2004), imply that PtdIns(4,5)*P*_2_ associates with inactive TOP2A in DNA-poor regions of the nucleus while PtdIns5*P* associates with active TOP2A on the chromatin. Depending on its activity and nuclear context, TOP2A may be associated with PtdIns(4,5)*P*_2_ or PtdIns5*P*. PtdIns5*P* binds to other nuclear proteins and their interaction can have an impact on their chromatin association and functional activity. For example, PtdIns5*P* binds to the histone code reader inhibitor of growth protein 2 (ING2) via a PBR and this interaction has been described to promote chromatin association to specific gene promoters involved in the DNA damage response, via the inhibition of PIP5K2B, (Gozani et al., 2003; Jones et al., 2006; Bua et al., 2013). In contrast, PtdIns5*P* binds to sin3A-associated protein 30 (SAP30) and this interaction led to the dissociation of SAP30 from DNA, which negatively impacted transcription (Viiri et al., 2009).

TOP2A has been described to also localise to the nucleolus in interphase (Meyer et al., 1997; Agostinho et al., 2004; Ray et al., 2013) but this localisation is more prominent when it is overexpressed (Christensen et al., 2002; Tavormina et al., 2002; Agostinho et al., 2004; Linka et al., 2007; Morotomi-Yano & Yano, 2021). Overexpression of the full length or the CTD of TOP2A in HeLa cells led to its accumulation in the nucleolus, consistently with the aforementioned studies. We have now identified two regions necessary for the nucleolar localisation of TOP2A, consisting of the PBR, M2 (1228-1237) and M3-4 (1259-1276), located in the proximal region of the CTD. The location of these regions is consistent with the NoLS identified in the rat TOP2A orthologue, spanning the residues 1192-1289 (Yasuda et al., 2021), and which corresponds to residues 1195-1293 in human TOP2A, hence including both M2 and M3-4 in the human enzyme. The NoLS identified in the rat orthologue was also shown to interact with RNA and this molecular association kept TOP2A in an inactive state (Yasuda et al., 2021). The nucleolar localisation of TOP2A is also observed endogenously in HeLa cells but only in about five percent of fixed cells from an actively growing population (data not shown). It was shown to accumulate in this sub-nuclear compartment upon ATP depletion and its nucleolar retention was dependent upon rRNA transcription (Morotomi-Yano & Yano, 2021). In addition, the salt-insoluble fraction of TOP2A was revealed to locate in the nucleolus and described as inactive (Agostinho et al., 2004). Overall, the two PBR act as nucleolar localisation sequences, and as interaction sites to RNA and PPIn and may retain TOP2A in an inactive state. rRNA was shown to be necessary for the retention of TOP2A in the nucleolus (Morotomi-Yano & Yano, 2021) but how PPIn could contribute is still unclear. Other PPIn-interacting proteins were shown to harbour PBR with dual function in both PPIn interaction and nucleolar localisation, such as proliferation-associated 2G4 (PA2G4 or EBP1) and syntenin-2 (Mortier et al., 2005; Karlsson et al., 2016), suggesting a common mechanism for these proteins. Substitution of lysine to uncharged residues in the PPIn-binding PBR resulted both in loss of PPIn binding and exclusion from the nucleolus (Mortier et al., 2005; Karlsson et al., 2016).

In summary, two PBR located in the proximal part of the CTD of TOP2A act as sequences contributing to nucleolar localisation and PPIn interaction. In line with the study from Yasuda *et al* on the rat orthologue, the two PBR in the human enzyme are mapped in the same region of the rat enzyme that keeps it in an inactive state in nucleoli via its association with RNA (Yasuda et al., 2021), potentially rRNA (Morotomi-Yano & Yano, 2021). We would therefore speculate that the 2 PBR in the human enzyme act in the same way. Nuclear speckles may also act as a reservoir of inactive TOP2A in association with PtdIns(4,5)*P*_2_, but how TOP2A is targeted to this site is unclear. A previous study by Lane *et al* showed that another polybasic cluster located in the distal part of the CTD (residues 1501-1531) is necessary for TOP2A to anchor to mitotic chromatin (Lane et al., 2013). Overall, these studies indicate that different parts of the CTD may allow TOP2A to discriminate between DNA and RNA binding defining different functions and subcellular localisation for the enzyme. How PPIn interaction contributes to these properties remains to be clarified.

## Acknowledgements

We thank Susan PC Cole (Queen’s University, Canada) and William Beck (University of Illinois, USA) for kind gifts of plasmids. This work was funded by L. Meltzers Høyskolefond, Nansenfondet and the University of Bergen (likestilingsmidler).

## References

Agostinho, M., Rino, J., Braga, J., Ferreira, F., Steffensen, S., & Ferreira, J. (2004). Human topoisomerase IIalpha: targeting to subchromosomal sites of activity during interphase and mitosis. Molecular biology of the cell, 15(5), 2388–2400. doi:10.1091/mbc.e03-08-0558

Ahn, J. Y., Liu, X., Cheng, D., Peng, J., Chan, P. K., Wade, P. A., & Ye, K. (2005). Nucleophosmin/B23, a nuclear PI(3,4,5)P(3) receptor, mediates the antiapoptotic actions of NGF by inhibiting CAD. Mol Cell, 18(4), 435–445. doi:10.1016/j.molcel.2005.04.010

Albi, E., Cataldi, S., Rossi, G., & Magni, M. V. (2003). A possible role of cholesterol-sphingomyelin/phosphatidylcholine in nuclear matrix during rat liver regeneration. J Hepatol, 38(5), 623–628. doi:S0168827803000746 [pii]

Bayona-Hernandez, A., Guerra, S., Jimenez-Ramirez, I. A., Sztacho, M., Hozak, P., Rodriguez-Zapata, L. C., Pereira-Santana, A., & Castano, E. (2023). LIPRNAseq: a method to discover lipid interacting RNAs by sequencing. Mol Biol Rep, 50(8), 6619–6626. doi:10.1007/s11033-023-08548-5

Bidlingmaier, S., Wang, Y., Liu, Y., Zhang, N., & Liu, B. (2011). Comprehensive analysis of yeast surface displayed cDNA library selection outputs by exon microarray to identify novel protein-ligand interactions. Mol Cell Proteomics, 10(3), M110 005116. doi:10.1074/mcp.M110.005116

Blind, R. D., Sablin, E. P., Kuchenbecker, K. M., Chiu, H. J., Deacon, A. M., Das, D., Fletterick, R. J., & Ingraham, H. A. (2014). The signaling phospholipid PIP3 creates a new interaction surface on the nuclear receptor SF-1. Proc Natl Acad Sci U S A, 111(42), 15054–15059. doi:10.1073/pnas.1416740111

Blind, R. D., Suzawa, M., & Ingraham, H. A. (2012). Direct Modification and Activation of a Nuclear Receptor-PIP2 Complex by the Inositol Lipid Kinase IPMK. Sci Signal, 5(229). doi:ARTN ra44 DOI 10.1126/scisignal.2003111

Boege, F., Andersen, A., Jensen, S., Zeidler, R., & Kreipe, H. (1995). Proliferation-associated nuclear antigen Ki-S1 is identical with topoisomerase II alpha. Delineation of a carboxy-terminal epitope with peptide antibodies. Am J Pathol, 146(6), 1302–1308. Retrieved from https://www.ncbi.nlm.nih.gov/pubmed/7539979

Boronenkov, I. V., Loijens, J. C., Umeda, M., & Anderson, R. A. (1998). Phosphoinositide signaling pathways in nuclei are associated with nuclear speckles containing pre-mRNA processing factors. Mol Biol Cell, 9(12), 3547–3560. doi:10.1091/mbc.9.12.3547

Bua, D. J., Martin, G. M., Binda, O., & Gozani, O. (2013). Nuclear phosphatidylinositol-5-phosphate regulates ING2 stability at discrete chromatin targets in response to DNA damage. Sci Rep, 3, 2137. doi:10.1038/srep02137

Bultsma, Y., Keune, W. J., & Divecha, N. (2010). PIP4Kbeta interacts with and modulates nuclear localization of the high-activity PtdIns5P-4-kinase isoform PIP4Kalpha. Biochem J, 430(2), 223–235. doi:10.1042/BJ20100341

Chen, C., Chen, M., Wen, T., Anderson, R. A., & Cryns, V. L. (2023). Regulation of NRF2 by Phosphoinositides and Small Heat Shock Proteins. bioRxiv. doi:10.1101/2023.10.26.564194

Choi, S., Chen, M., Cryns, V. L., & Anderson, R. A. (2019). A nuclear phosphoinositide kinase complex regulates p53. Nat Cell Biol, 21(4), 462–475. doi:10.1038/s41556-019-0297-2

Christensen, M. O., Larsen, M. K., Barthelmes, H. U., Hock, R., Andersen, C. L., Kjeldsen, E., Knudsen, B. R., Westergaard, O., Boege, F., & Mielke, C. (2002). Dynamics of human DNA topoisomerases IIalpha and IIbeta in living cells. The Journal of cell biology, 157(1), 31–44. doi:10.1083/jcb.200112023

Ciruela, A., Hinchliffe, K. A., Divecha, N., & Irvine, R. F. (2000). Nuclear targeting of the beta isoform of type II phosphatidylinositol phosphate kinase (phosphatidylinositol 5-phosphate 4-kinase) by its alpha-helix 7. Biochem J, 346 Pt 3(Pt 3), 587–591. Retrieved from https://www.ncbi.nlm.nih.gov/pubmed/10698683

Clarke, J. H., Letcher, A. J., D’Santos C, S., Halstead, J. R., Irvine, R. F., & Divecha, N. (2001). Inositol lipids are regulated during cell cycle progression in the nuclei of murine erythroleukaemia cells. Biochem J, 357(Pt 3), 905–910. doi:10.1042/0264-6021:3570905

Cocco, L., Gilmour, R. S., Ognibene, A., Letcher, A. J., Manzoli, F. A., & Irvine, R. F. (1987). Synthesis of polyphosphoinositides in nuclei of Friend cells. Evidence for polyphosphoinositide metabolism inside the nucleus which changes with cell differentiation. Biochem J, 248(3), 765–770. doi:10.1042/bj2480765

Di Lello, P., Nguyen, B. D., Jones, T. N., Potempa, K., Kobor, M. S., Legault, P., & Omichinski, J. G. (2005). NMR structure of the amino-terminal domain from the Tfb1 subunit of TFIIH and characterization of its phosphoinositide and VP16 binding sites. Biochemistry, 44(21), 7678–7686. doi:10.1021/bi050099s

Divecha, N., Banfic, H., & Irvine, R. F. (1991). The polyphosphoinositide cycle exists in the nuclei of Swiss 3T3 cells under the control of a receptor (for IGF-I) in the plasma membrane, and stimulation of the cycle increases nuclear diacylglycerol and apparently induces translocation of protein kinase C to the nucleus. EMBO J, 10(11), 3207–3214. doi:10.1002/j.1460-2075.1991.tb04883.x

Drake, F. H., Hofmann, G. A., Bartus, H. F., Mattern, M. R., Crooke, S. T., & Mirabelli, C. K. (1989). Biochemical and pharmacological properties of p170 and p180 forms of topoisomerase II. Biochemistry, 28(20), 8154–8160. doi:10.1021/bi00446a029

Elong Edimo, W., Derua, R., Janssens, V., Nakamura, T., Vanderwinden, J. M., Waelkens, E., & Erneux, C. (2011). Evidence of SHIP2 Ser132 phosphorylation, its nuclear localization and stability. Biochem J, 439(3), 391–401. doi:10.1042/BJ20110173

Endsley, C. E., Moore, K. A., Townsley, T. D., Durston, K. K., & Deweese, J. E. (2024). Bioinformatic Analysis of Topoisomerase IIalpha Reveals Interdomain Interdependencies and Critical C-Terminal Domain Residues. Int J Mol Sci, 25(11). doi:10.3390/ijms25115674

Fujimoto, T. (2024). Nuclear lipid droplet: Guardian of nuclear membrane lipid homeostasis? Curr Opin Cell Biol, 88, 102370. doi:10.1016/j.ceb.2024.102370

Galganski, L., Urbanek, M. O., & Krzyzosiak, W. J. (2017). Nuclear speckles: molecular organization, biological function and role in disease. Nucleic Acids Res, 45(18), 10350–10368. doi:10.1093/nar/gkx759

Gardiner, L. P., Roper, D. I., Hammonds, T. R., & Maxwell, A. (1998). The N-terminal domain of human topoisomerase IIalpha is a DNA-dependent ATPase. Biochemistry, 37(48), 16997–17004. doi:10.1021/bi9818321

Gelato, K. A., Tauber, M., Ong, M. S., Winter, S., Hiragami-Hamada, K., Sindlinger, J., Lemak, A., Bultsma, Y., Houliston, S., Schwarzer, D., Divecha, N., Arrowsmith, C. H., & Fischle, W. (2014). Accessibility of different histone H3-binding domains of UHRF1 is allosterically regulated by phosphatidylinositol 5-phosphate. Mol Cell, 54(6), 905–919. doi:10.1016/j.molcel.2014.04.004

Gillooly, D. J., Morrow, I. C., Lindsay, M., Gould, R., Bryant, N. J., Gaullier, J. M., Parton, R. G., & Stenmark, H. (2000). Localization of phosphatidylinositol 3-phosphate in yeast and mammalian cells. EMBO J, 19(17), 4577–4588. doi:10.1093/emboj/19.17.4577

Goswami, P. C., Roti Roti, J. L., & Hunt, C. R. (1996). The cell cycle-coupled expression of topoisomerase IIalpha during S phase is regulated by mRNA stability and is disrupted by heat shock or ionizing radiation. Molecular and cellular biology, 16(4), 1500–1508. doi:10.1128/MCB.16.4.1500

Gozani, O., Karuman, P., Jones, D. R., Ivanov, D., Cha, J., Lugovskoy, A. A., Baird, C. L., Zhu, H., Field, S. J., Lessnick, S. L., Villasenor, J., Mehrotra, B., Chen, J., Rao, V. R., Brugge, J. S., Ferguson, C. G., Payrastre, B., Myszka, D. G., Cantley, L. C., Wagner, G., Divecha, N., Prestwich, G. D., & Yuan, J. (2003). The PHD finger of the chromatin-associated protein ING2 functions as a nuclear phosphoinositide receptor. Cell, 114(1), 99–111. doi:10.1016/s0092-8674(03)00480-x

Huang, W., Zhang, H., Davrazou, F., Kutateladze, T. G., Shi, X., Gozani, O., & Prestwich, G. D. (2007). Stabilized phosphatidylinositol-5-phosphate analogues as ligands for the nuclear protein ING2: chemistry, biology, and molecular modeling. J Am Chem Soc, 129(20), 6498–6506. doi:10.1021/ja070195b

Hunt, A. N. (2006). Dynamic lipidomics of the nucleus. J Cell Biochem, 97(2), 244–251. doi:10.1002/jcb.20691

Jacobsen, R. G., Mazloumi Gavgani, F., Edson, A. J., Goris, M., Altankhuyag, A., & Lewis, A. E. (2019). Polyphosphoinositides in the nucleus: Roadmap of their effectors and mechanisms of interaction. Adv Biol Regul, 72, 7–21. doi:10.1016/j.jbior.2019.04.001

Jenkins, J. R., Ayton, P., Jones, T., Davies, S. L., Simmons, D. L., Harris, A. L., Sheer, D., & Hickson, I. D. (1992). Isolation of cDNA clones encoding the beta isozyme of human DNA topoisomerase II and localisation of the gene to chromosome 3p24. Nucleic acids research, 20(21), 5587–5592. doi:10.1093/nar/20.21.5587

Jones, D. R., Bultsma, Y., Keune, W. J., Halstead, J. R., Elouarrat, D., Mohammed, S., Heck, A. J., D’Santos, C. S., & Divecha, N. (2006). Nuclear PtdIns5P as a transducer of stress signaling: an in vivo role for PIP4Kbeta. Mol Cell, 23(5), 685–695. doi:10.1016/j.molcel.2006.07.014

Kaadige, M. R., & Ayer, D. E. (2006). The polybasic region that follows the plant homeodomain zinc finger 1 of Pf1 is necessary and sufficient for specific phosphoinositide binding. J Biol Chem, 281(39), 28831–28836. doi:10.1074/jbc.M605624200

Kalasova, I., Faberova, V., Kalendova, A., Yildirim, S., Ulicna, L., Venit, T., & Hozak, P. (2016). Tools for visualization of phosphoinositides in the cell nucleus. Histochem Cell Biol, 145(4), 485–496. doi:10.1007/s00418-016-1409-8

Karlsson, T., Altankhuyag, A., Dobrovolska, O., Turcu, D. C., & Lewis, A. E. (2016). A polybasic motif in ErbB3-binding protein 1 (EBP1) has key functions in nucleolar localization and polyphosphoinositide interaction. Biochem J, 473(14), 2033–2047. doi:10.1042/BCJ20160274

Kitagawa, M., Liao, P. J., Lee, K. H., Wong, J., Shang, S. C., Minami, N., Sampetrean, O., Saya, H., Lingyun, D., Prabhu, N., Diam, G. K., Sobota, R., Larsson, A., Nordlund, P., McCormick, F., Ghosh, S., Epstein, D. M., Dymock, B. W., & Lee, S. H. (2017). Dual blockade of the lipid kinase PIP4Ks and mitotic pathways leads to cancer-selective lethality. Nat Commun, 8(1), 2200. doi:10.1038/s41467-017-02287-5

Krylova, I. N., Sablin, E. P., Moore, J., Xu, R. X., Waitt, G. M., MacKay, J. A., Juzumiene, D., Bynum, J. M., Madauss, K., Montana, V., Lebedeva, L., Suzawa, M., Williams, J. D., Williams, S. P., Guy, R. K., Thornton, J. W., Fletterick, R. J., Willson, T. M., & Ingraham, H. A. (2005). Structural analyses reveal phosphatidyl inositols as ligands for the NR5 orphan receptors SF-1 and LRH-1. Cell, 120(3), 343–355. doi:10.1016/j.cell.2005.01.024

Kwon, I. S., Lee, K. H., Choi, J. W., & Ahn, J. Y. (2010). PI(3,4,5)P3 regulates the interaction between Akt and B23 in the nucleus. BMB Rep, 43(2), 127–132. doi:10.5483/bmbrep.2010.43.2.127

Lane, A. B., Gimenez-Abian, J. F., & Clarke, D. J. (2013). A novel chromatin tether domain controls topoisomerase IIalpha dynamics and mitotic chromosome formation. The Journal of cell biology, 203(3), 471–486. doi:10.1083/jcb.201303045

Layerenza, J. P., Gonzalez, P., Garcia de Bravo, M. M., Polo, M. P., Sisti, M. S., & Ves-Losada, A. (2013). Nuclear lipid droplets: a novel nuclear domain. Biochim Biophys Acta, 1831(2), 327–340. doi:10.1016/j.bbalip.2012.10.005

Lewis, A. E., Sommer, L., Arntzen, M. O., Strahm, Y., Morrice, N. A., Divecha, N., & D’Santos, C. S. (2011). Identification of nuclear phosphatidylinositol 4,5-bisphosphate-interacting proteins by neomycin extraction. Mol Cell Proteomics, 10(2), M110 003376. doi:M110.003376 [pii] 10.1074/mcp.M110.003376

Lin, A., Hu, Q., Li, C., Xing, Z., Ma, G., Wang, C., Li, J., Ye, Y., Yao, J., Liang, K., Wang, S., Park, P. K., Marks, J. R., Zhou, Y., Zhou, J., Hung, M. C., Liang, H., Hu, Z., Shen, H., Hawke, D. H., Han, L., Zhou, Y., Lin, C., & Yang, L. (2017). The LINK-A lncRNA interacts with PtdIns(3,4,5)P(3) to hyperactivate AKT and confer resistance to AKT inhibitors. Nat Cell Biol, 19(3), 238–251. doi:10.1038/ncb3473

Lindsay, Y., McCoull, D., Davidson, L., Leslie, N., Fairservice, A., Gray, A., Lucocq, J., & Downes, C. (2006). Localization of agonist-sensitive PtdIns(3,4,5)P3 reveals a nuclear pool that is insensitive to PTEN expression. Journal of Cell Science, 119(Pt 24), 5160–5168. doi:10.1242/jcs.000133

Linka, R. M., Porter, A. C., Volkov, A., Mielke, C., Boege, F., & Christensen, M. O. (2007). C-terminal regions of topoisomerase IIalpha and IIbeta determine isoform-specific functioning of the enzymes in vivo. Nucleic acids research, 35(11), 3810–3822. doi:10.1093/nar/gkm102

Martin, T. F. (1998). Phosphoinositide lipids as signaling molecules: common themes for signal transduction, cytoskeletal regulation, and membrane trafficking. Annu Rev Cell Dev Biol, 14, 231–264. doi:10.1146/annurev.cellbio.14.1.231

Mate, S. M., Brenner, R. R., & Ves-Losada, A. (2006). Endonuclear lipids in liver cells. Can J Physiol Pharmacol, 84(3-4), 459–468. doi:10.1139/y05-097

Mazloumi Gavgani, F., Slinning, M. S., Morovicz, A. P., Arnesen, V. S., Turcu, D. C., Ninzima, S., D’Santos, C. S., & Lewis, A. E. (2021). Nuclear Phosphatidylinositol 3,4,5-Trisphosphate Interactome Uncovers an Enrichment in Nucleolar Proteins. Mol Cell Proteomics, 20, 100102. doi:10.1016/j.mcpro.2021.100102

Mazzotti, G., Zini, N., Rizzi, E., Rizzoli, R., Galanzi, A., Ognibene, A., Santi, S., Matteucci, A., Martelli, A. M., & Maraldi, N. M. (1995). Immunocytochemical detection of phosphatidylinositol 4,5-bisphosphate localization sites within the nucleus. J Histochem Cytochem, 43(2), 181–191. doi:10.1177/43.2.7822774

McClendon, A. K., Gentry, A. C., Dickey, J. S., Brinch, M., Bendsen, S., Andersen, A. H., & Osheroff, N. (2008). Bimodal recognition of DNA geometry by human topoisomerase II alpha: preferential relaxation of positively supercoiled DNA requires elements in the C-terminal domain. Biochemistry, 47(50), 13169–13178. doi:10.1021/bi800453h

McClendon, A. K., & Osheroff, N. (2007). DNA topoisomerase II, genotoxicity, and cancer. Mutation research, 623(1-2), 83–97. doi:10.1016/j.mrfmmm.2007.06.009

McClendon, A. K., Rodriguez, A. C., & Osheroff, N. (2005). Human topoisomerase IIalpha rapidly relaxes positively supercoiled DNA: implications for enzyme action ahead of replication forks. The Journal of biological chemistry, 280(47), 39337–39345. doi:10.1074/jbc.M503320200

McPhee, M. J., Salsman, J., Foster, J., Thompson, J., Mathavarajah, S., Dellaire, G., & Ridgway, N. D. (2022). Running ’LAPS’ Around nLD: Nuclear Lipid Droplet Form and Function. Front Cell Dev Biol, 10, 837406. doi:10.3389/fcell.2022.837406

Meczes, E. L., Gilroy, K. L., West, K. L., & Austin, C. A. (2008). The impact of the human DNA topoisomerase II C-terminal domain on activity. PloS one, 3(3), e1754. doi:10.1371/journal.pone.0001754

Mellman, D. L., Gonzales, M. L., Song, C., Barlow, C. A., Wang, P., Kendziorski, C., & Anderson, R. A. (2008). A PtdIns4,5P2-regulated nuclear poly(A) polymerase controls expression of select mRNAs. Nature, 451(7181), 1013–1017. doi:nature06666 [pii] 10.1038/nature06666

Mendez, J., & Stillman, B. (2000). Chromatin association of human origin recognition complex, cdc6, and minichromosome maintenance proteins during the cell cycle: assembly of prereplication complexes in late mitosis. Mol Cell Biol, 20(22), 8602–8612. doi:10.1128/MCB.20.22.8602-8612.2000

Meyer, K. N., Kjeldsen, E., Straub, T., Knudsen, B. R., Hickson, I. D., Kikuchi, A., Kreipe, H., & Boege, F. (1997). Cell cycle-coupled relocation of types I and II topoisomerases and modulation of catalytic enzyme activities. The Journal of cell biology, 136(4), 775–788. doi:10.1083/jcb.136.4.775

Michell, R. H., Heath, V. L., Lemmon, M. A., & Dove, S. K. (2006). Phosphatidylinositol 3,5-bisphosphate: metabolism and cellular functions. Trends Biochem Sci, 31(1), 52–63. doi:S0968-0004(05)00343-9 [pii] 10.1016/j.tibs.2005.11.013

Miladinovic, A., Antiga, L., Venit, T., Bayona-Hernandez, A., Cervenka, J., Labala, R. K., Kolar, M., Castano, E., Sztacho, M., & Hozak, P. (2024). The perinucleolar compartment and the oncogenic super-enhancers are part of the same phase-separated structure filled with phosphatidylinositol 4,5bisphosphate and long noncoding RNA HANR. Adv Biol Regul, 101069. doi:10.1016/j.jbior.2024.101069

Mirski, S. E., Gerlach, J. H., & Cole, S. P. (1999). Sequence determinants of nuclear localization in the alpha and beta isoforms of human topoisomerase II. Exp Cell Res, 251(2), 329–339. doi:10.1006/excr.1999.4587

Morotomi-Yano, K., & Yano, K. I. (2021). Nucleolar translocation of human DNA topoisomerase II by ATP depletion and its disruption by the RNA polymerase I inhibitor BMH-21. Sci Rep, 11(1), 21533. doi:10.1038/s41598-021-00958-4

Morovicz, A. P., Mazloumi Gavgani, F., Jacobsen, R. G., Skuseth Slinning, M., Turcu, D. C., & Lewis, A. E. (2022). Phosphoinositide 3-kinase signalling in the nucleolus. Adv Biol Regul, 83, 100843. doi:10.1016/j.jbior.2021.100843

Mortier, E., Wuytens, G., Leenaerts, I., Hannes, F., Heung, M. Y., Degeest, G., David, G., & Zimmermann, P. (2005). Nuclear speckles and nucleoli targeting by PIP2-PDZ domain interactions. EMBO J, 24(14), 2556–2565. doi:10.1038/sj.emboj.7600722

Okada, M., Jang, S. W., & Ye, K. (2008). Akt phosphorylation and nuclear phosphoinositide association mediate mRNA export and cell proliferation activities by ALY. Proc Natl Acad Sci U S A, 105(25), 8649–8654. doi:0802533105 [pii] 10.1073/pnas.0802533105

Osborne, S. L., Thomas, C. L., Gschmeissner, S., & Schiavo, G. (2001). Nuclear PtdIns(4,5)P2 assembles in a mitotically regulated particle involved in pre-mRNA splicing. J Cell Sci, 114(Pt 13), 2501–2511. doi:10.1242/jcs.114.13.2501

Park, S. W., Parrott, A. M., Fritz, D. T., Park, Y., Mathews, M. B., & Lee, C. G. (2008). Regulation of the catalytic function of topoisomerase II alpha through association with RNA. Nucleic acids research, 36(19), 6080–6090. doi:10.1093/nar/gkn614

Pommier, Y., Nussenzweig, A., Takeda, S., & Austin, C. (2022). Human topoisomerases and their roles in genome stability and organization. Nat Rev Mol Cell Biol, 23(6), 407–427. doi:10.1038/s41580-022-00452-3

Postle, A. D., Wilton, D. C., Hunt, A. N., & Attard, G. S. (2007). Probing phospholipid dynamics by electrospray ionisation mass spectrometry. Prog Lipid Res, 46(3-4), 200–224. doi:S0163-7827(07)00007-0 [pii] 10.1016/j.plipres.2007.04.001

Rattner, J. B., Hendzel, M. J., Furbee, C. S., Muller, M. T., & Bazett-Jones, D. P. (1996). Topoisomerase II alpha is associated with the mammalian centromere in a cell cycle- and species-specific manner and is required for proper centromere/kinetochore structure. The Journal of cell biology, 134(5), 1097–1107. doi:10.1083/jcb.134.5.1097

Ray, S., Panova, T., Miller, G., Volkov, A., Porter, A. C., Russell, J., Panov, K. I., & Zomerdijk, J. C. (2013). Topoisomerase IIalpha promotes activation of RNA polymerase I transcription by facilitating pre-initiation complex formation. Nat Commun, 4, 1598. doi:10.1038/ncomms2599

Sablin, E. P., Blind, R. D., Uthayaruban, R., Chiu, H. J., Deacon, A. M., Das, D., Ingraham, H. A., & Fletterick, R. J. (2015). Structure of Liver Receptor Homolog-1 (NR5A2) with PIP3 hormone bound in the ligand binding pocket. J Struct Biol, 192(3), 342–348. doi:10.1016/j.jsb.2015.09.012

Samardak, K., Bacle, J., & Moriel-Carretero, M. (2024). Behind the stoNE wall: A fervent activity for nuclear lipids. Biochimie. doi:10.1016/j.biochi.2024.08.002

Sarkes, D., & Rameh, L. E. (2010). A novel HPLC-based approach makes possible the spatial characterization of cellular PtdIns5P and other phosphoinositides. Biochemical Journal, 428, 375–384. doi:10.1042/Bj20100129

Sarwar, M., Syed Khaja, A. S., Aleskandarany, M., Karlsson, R., Althobiti, M., Odum, N., Mongan, N. P., Dizeyi, N., Johnson, H., Green, A. R., Ellis, I. O., Rakha, E. A., & Persson, J. L. (2019). The role of PIP5K1alpha/pAKT and targeted inhibition of growth of subtypes of breast cancer using PIP5K1alpha inhibitor. Oncogene, 38(3), 375–389. doi:10.1038/s41388-018-0438-2

Schindelin, J., Arganda-Carreras, I., Frise, E., Kaynig, V., Longair, M., Pietzsch, T., Preibisch, S., Rueden, C., Saalfeld, S., Schmid, B., Tinevez, J. Y., White, D. J., Hartenstein, V., Eliceiri, K., Tomancak, P., & Cardona, A. (2012). Fiji: an open-source platform for biological-image analysis. Nat Methods, 9(7), 676–682. doi:10.1038/nmeth.2019

Scott, M. S., Troshin, P. V., & Barton, G. J. (2011). NoD: a Nucleolar localization sequence detector for eukaryotic and viral proteins. BMC Bioinformatics, 12, 317. doi:10.1186/1471-2105-12-317

Semenas, J., Hedblom, A., Miftakhova, R. R., Sarwar, M., Larsson, R., Shcherbina, L., Johansson, M. E., Harkonen, P., Sterner, O., & Persson, J. L. (2014). The role of PI3K/AKT-related PIP5K1alpha and the discovery of its selective inhibitor for treatment of advanced prostate cancer. Proc Natl Acad Sci U S A, 111(35), E3689–3698. doi:10.1073/pnas.1405801111

Skare, P., & Karlsson, R. (2002). Evidence for two interaction regions for phosphatidylinositol(4,5)-bisphosphate on mammalian profilin I. FEBS Lett, 522(1-3), 119–124. doi:10.1016/s0014-5793(02)02913-7

Sobol, M., Krausova, A., Yildirim, S., Kalasova, I., Faberova, V., Vrkoslav, V., Philimonenko, V., Marasek, P., Pastorek, L., Capek, M., Lubovska, Z., Ulicna, L., Tsuji, T., Lisa, M., Cvacka, J., Fujimoto, T., & Hozak, P. (2018). Nuclear phosphatidylinositol 4,5-bisphosphate islets contribute to efficient RNA polymerase II-dependent transcription. J Cell Sci, 131(8). doi:10.1242/jcs.211094

Sosa Ponce, M. L., Cobb, J. A., & Zaremberg, V. (2024). Lipids and chromatin: a tale of intriguing connections shaping genomic landscapes. Trends Cell Biol. doi:10.1016/j.tcb.2024.06.004

Stijf-Bultsma, Y., Sommer, L., Tauber, M., Baalbaki, M., Giardoglou, P., Jones, D. R., Gelato, K. A., van Pelt, J., Shah, Z., Rahnamoun, H., Toma, C., Anderson, K. E., Hawkins, P., Lauberth, S. M., Haramis, A. P., Hart, D., Fischle, W., & Divecha, N. (2015). The basal transcription complex component TAF3 transduces changes in nuclear phosphoinositides into transcriptional output. Mol Cell, 58(3), 453–467. doi:10.1016/j.molcel.2015.03.009

Tabellini, G., Bortul, R., Santi, S., Riccio, M., Baldini, G., Cappellini, A., Billi, A. M., Berezney, R., Ruggeri, A., Cocco, L., & Martelli, A. M. (2003). Diacylglycerol kinase-theta is localized in the speckle domains of the nucleus. Exp Cell Res, 287(1), 143–154. doi:10.1016/s0014-4827(03)00115-0

Tavormina, P. A., Come, M. G., Hudson, J. R., Mo, Y. Y., Beck, W. T., & Gorbsky, G. J. (2002). Rapid exchange of mammalian topoisomerase II alpha at kinetochores and chromosome arms in mitosis. The Journal of cell biology, 158(1), 23–29. doi:10.1083/jcb.200202053

Toska, E., Campbell, H. A., Shandilya, J., Goodfellow, S. J., Shore, P., Medler, K. F., & Roberts, S. G. (2012). Repression of transcription by WT1-BASP1 requires the myristoylation of BASP1 and the PIP2-dependent recruitment of histone deacetylase. Cell Rep, 2(3), 462–469. doi:10.1016/j.celrep.2012.08.005

Vanden Broeck, A., Lotz, C., Drillien, R., Haas, L., Bedez, C., & Lamour, V. (2021). Structural basis for allosteric regulation of Human Topoisomerase IIalpha. Nat Commun, 12(1), 2962. doi:10.1038/s41467-021-23136-6

Vann, L. R., Wooding, F. B., Irvine, R. F., & Divecha, N. (1997). Metabolism and possible compartmentalization of inositol lipids in isolated rat-liver nuclei. Biochem J, 327 (Pt 2)(Pt 2), 569–576. doi:10.1042/bj3270569

Vidalle, M. C., Sheth, B., Fazio, A., Marvi, M. V., Leto, S., Koufi, F. D., Neri, I., Casalin, I., Ramazzotti, G., Follo, M. Y., Ratti, S., Manzoli, L., Gehlot, S., Divecha, N., & Fiume, R. (2023). Nuclear Phosphoinositides as Key Determinants of Nuclear Functions. Biomolecules, 13(7). doi:10.3390/biom13071049

Viiri, K. M., Janis, J., Siggers, T., Heinonen, T. Y., Valjakka, J., Bulyk, M. L., Maki, M., & Lohi, O. (2009). DNA-binding and -bending activities of SAP30L and SAP30 are mediated by a zinc-dependent module and monophosphoinositides. Mol Cell Biol, 29(2), 342–356. doi:MCB.01213-08 [pii] 10.1128/MCB.01213-08

Vos, S., Tretter, E., Schmidt, B., & Berger, J. (2011). All tangled up: how cells direct, manage and exploit topoisomerase function. Nature reviews. Molecular cell biology, 12(12), 827–841. doi:10.1038/nrm3228

Wang, Y. H., & Sheetz, M. P. (2022). When PIP(2) Meets p53: Nuclear Phosphoinositide Signaling in the DNA Damage Response. Front Cell Dev Biol, 10, 903994. doi:10.3389/fcell.2022.903994

Wang, Y. R., Chen, S. F., Wu, C. C., Liao, Y. W., Lin, T. S., Liu, K. T., Chen, Y. S., Li, T. K., Chien, T. C., & Chan, N. L. (2017). Producing irreversible topoisomerase II-mediated DNA breaks by site-specific Pt(II)-methionine coordination chemistry. Nucleic Acids Res, 45(18), 10861–10871. doi:10.1093/nar/gkx742

Watt, S. A., Kular, G., Fleming, I. N., Downes, C. P., & Lucocq, J. M. (2002). Subcellular localization of phosphatidylinositol 4,5-bisphosphate using the pleckstrin homology domain of phospholipase C delta1. Biochem J, 363(Pt 3), 657–666. doi:10.1042/0264-6021:3630657

Wendorff, T. J., Schmidt, B. H., Heslop, P., Austin, C. A., & Berger, J. M. (2012). The structure of DNA-bound human topoisomerase II alpha: conformational mechanisms for coordinating inter-subunit interactions with DNA cleavage. J Mol Biol, 424(3-4), 109–124. doi:10.1016/j.jmb.2012.07.014

Woessner, R. D., Mattern, M. R., Mirabelli, C. K., Johnson, R. K., & Drake, F. H. (1991). Proliferation- and cell cycle-dependent differences in expression of the 170 kilodalton and 180 kilodalton forms of topoisomerase II in NIH-3T3 cells. Cell growth & differentiation : the molecular biology journal of the American Association for Cancer Research, 2(4), 209–214. Retrieved from https://www.ncbi.nlm.nih.gov/pubmed/1651102

Yasuda, K., Kato, Y., Ikeda, S., & Kawano, S. (2021). Regulation of catalytic activity and nucleolar localization of rat DNA topoisomerase IIalpha through its C-terminal domain. Genes Genet Syst, 95(6), 291–302. doi:10.1266/ggs.20-00038

Yildirim, S., Castano, E., Sobol, M., Philimonenko, V. V., Dzijak, R., Venit, T., & Hozak, P. (2013). Involvement of phosphatidylinositol 4,5-bisphosphate in RNA polymerase I transcription. J Cell Sci, 126(Pt 12), 2730–2739. doi:10.1242/jcs.123661

York, J. D., & Majerus, P. W. (1994). Nuclear phosphatidylinositols decrease during S-phase of the cell cycle in HeLa cells. J Biol Chem, 269(11), 7847–7850. Retrieved from https://www.ncbi.nlm.nih.gov/pubmed/8132500

