## Supplemental figures for "Lysine-rich regions located in the C-terminal domain of DNA Topoisomerase 2-alpha act as polyphosphoinositide interaction sites and nucleolar localisation signals"

#### Supplementary Figure S1

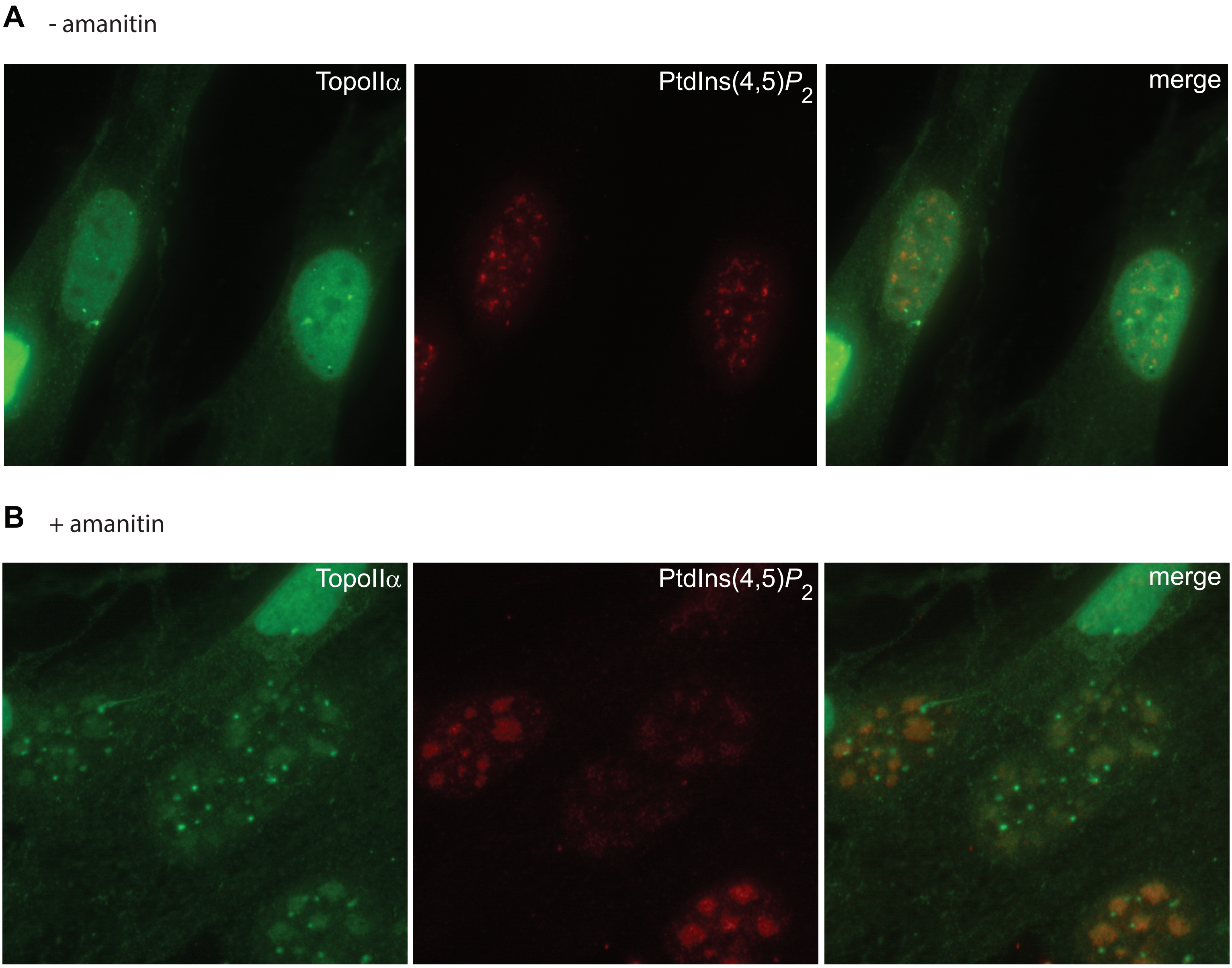


**Supplementary Figure S1:**
**Endogenous Topo IIα colocalises with PtdIns(4,5)*P*2 in nuclear speckles upon RNA Pol II inhibition in TIG-1 cells**
**A,** Asynchronous and **B,** α-amanitin (50 μg/mL, 5h) treated TIG-1 cells stained with the anti-Topo IIα KiS1 monoclonal antibody and anti-PtdIns(4,5)P2 IgM antibody; white squares denote nuclei that are enlarged in the panel below. Scale bar, 10 μm.

### Supplementary Figure S2

```
                                               M1                                    M2
Human   1171 LATFIEELEAVEAKEKQDEQVGLPGKGGKAKG---------KKTQMA-EVLPSPRGQRVIPRITIEMKAEAEKKNKKKI
Mouse   1168 LAVFIEELEVVEAKEKQDEQVGLPGKAGKAKG---------KKAQMCADVLPSPRGKRVIPQVTVEMKAEAEKKIRKKI
Rat     1168 LAAFVEELEVVEAKEKQDEQVGLPGKGVKAKG---------KKAQIS-EVLPSPVGKRVIPQVTMEMRAEAEKKIRRKI
Pig     1171 LAAFIEELEAVEAKEKQDEQIGLPGKGGKAKG---------KKTQMA-EVLPSPCGKRVIPRVTVEMKAEAEKKIKKKI
Chicken 1163 LAAFVEELDAVEAKQMQDEMAGITGKPLKVKGGKQGGKQKVTKAQLA-EVMPSPHGIRVVPRVTAEMKAEAEKRIKKKI
Hamster 1166 LAVFIEELEVVEAKEKQDEQVGLPGKGGKAKG---------KKAQMS-EVLPSPHGKRVIPQVTMEMKAEAEKKIRKKI
             **.*:***:.****: ***  *: **  *.**         .*:*:. :*:*** * **:*::* **:*****: ::**

                                           M3-4                M5
Human   1240 KNENTEGSPQEDG-----VELEGLKQRLEKKQKREPGTKTKKQTTLAFKPIKKGKKRNPWSDSESDRSSDESNFDVPPR
Mouse   1238 KSENVEGTPAEDG-----AEPGSLRQRIEKKQKKEPGA--KKQTTLPFKPVKKGRKKNPWSDSESDVSSNESNVDVPPR
Rat     1237 KSENVEGTPAEDG-----AEPG-LRQRLEKRQKREPGTRAKKQTTLPFKPIKKAQKQNPWSDSESDMSSNESNFDVPPR
Pig     1240 KSENTEGSPQEDG-----MEVEGLKQRLEKKQKREPGTKTKKQTTLPFKPIKKAKKRNPWSDSESDISSDESNFNVPPR
Chicken 1241 KSEKNESDEKQEGNSSGDKEPSSLKQRLAQKRKAEQGT--KRQTTLPFKPIKKM-KRNPWSDSESDSESDD--FEVPSK
Hamster 1235 KSENVEGTPTENG-----LELGSLKQRIEKKQKKEPGAMTKKQTTLAFKPIKKGKKRNPWSDSESDMSSNESNVDVPPR
             *.*: *.   ::*      *   *:**: :::* * *:  *:**** ***:** *:********* .*::  .:** :

Human   1314 -ETEPRRAATKTKFTMDLDSDEDFSDFDEKTDDEDF--------VPSDASPPKTKTSPKLSNKELKPQKS--VVSDLEA
Mouse   1310 -QKEQRSAAAKAKFTVDLDSDEDFSGLDEKDEDEDF--------LPLDATPPKAKIPPKNTKKALKTQGSSMSVVDLES
Rat     1311 -EKEPRIAATKAKFTADLDSDDDFSGLDEKDEDEDF--------FPLDDTPPKTKMPPKNTKKALKPQKSSTS-VDLES
Pig     1314 -EKEPRRAAAKTKFTVDLDSDEDFSDADEKTRDEDF--------VPSDTSPQKAETSPKHTNKEPKPQKSTPSVSDFDA
Chicken 1315 RERVVRQAAAKIKPMVNSDSDADLTSSDEDSEYQENSEGNTDSDTTSKKKPPKAKAVPKEKKGKAPKEKPLPDAVPVRV
Hamster 1309 -EKDPRRAATKAKFTMDLDSDEDFSGSDGKDEDEDF--------FPLDTTPPKTKIPQKNTKKALKPQKSAMS-GDPES
              :   * **:* *   : *** *::. * .   ::            . .* *::   * .:     :

Human   1382 DD-VKGSVPLSSSPPATHFPDETEITNPVPKKNVTVKK--TAAKSQSSTSTTGAKKRAAPKGTKR-------DPALNSG
Mouse   1380 -D-VKDSVPASPGVPAADFPAETEQSKPS-KKTVGVKK--TATKSQSSVSTAGTKKRAAPKGTKS-------DSALSAR
Rat     1379 -D-GKDSVPASPGASAADVPAETEPSKPSSKQTVGVKR--TITKGQSLTSTAGTKKRAVPKETKS-------DSALNAH
Pig     1384 DD-AKDNVPPSPSSPVADFPAVTETIKPVSKKNVTVKK--TAAKSQSSTSTTGAKKRAAPKGAKK-------DPDLDSD
Chicken 1394 QNVAAESASQDPAAPPVSVPRAQ----AVPKKPAAAKKGSTAKDNQPSIMDILTKKKAAPKAPRRAQREESPPSEATAA
Hamster 1378 -D-EKDSVPASPGPPAADLPADTEQLKPSSKQTVAVKK--TATKSQSSTSTAGTKKRAVPKGSKS-------DSALNAH
              :    ..  . .   . .*          *: . .*:  *  ..*       :**:*.**  :             :

                    M6
Human   1451 VSQKPDPAKTKNRRKRKPSTSDDSDSNFEKIVSKAVTSKKSKGESD-DFH----MDFDSAVAPRAKSVRAKKPIKYLEE
Mouse   1447 VSEKPAPAKAKNSRKRKPSSSDSSDSDFERAISKGATSKKAKGEEQ-DFP----VDLEDTIAPRAKSDRARKPIKYLEE
Rat     1447 VSKKPAPAKAKNSRKRMPSSSDSSDSEFEKAISKGATSKKLKGEER-DFH----VDLDDTVAPRAKSGRARKPIKYLEE
Pig     1453 VSKKPNPPKPKGRRKRKPSTSDDSDSNFEKMISKAVTSKKPKGESD-DFH----LDLDLAVASRAKSGRTKKPIKYLEE
Chicken 1469 VAKKPGPPRGRKATKRLTS-SSDSDSDFGSRPSKSVAAKKSKRDDDDSYSIDLTADSPAAAAPRTRPGRLKKPVQYLES
Hamster 1466 GPEKPVPAKAKNSRKRKQSSSDDSDSDFEKVVSKVAASKKSKGENQ-DFR----VDLDETMVPRAKSGRAKKPIKYLEE
               :** * : :   **  * *..***:*    ** .::** * :.  .:      *   : . *::  * :**::***.

Human   1525 SDEDDLF- 1531
Mouse   1521 SDDDDDLF 1528
Rat     1521 SDDDLF-- 1526
Pig     1527 SDEDDLF- 1533
Chicken 1547 SDEDDMF- 1553
Hamster 1520 SDDDDLF- 1526
             **:*
```

**Supplementary Figure S2. Alignment of vertebrate TOP2A CTD.**
Sequences from Human (UniProtKB: P11388-1, residues 1171-1531), mouse (Mus musculus, UniProtKB: Q01320-1, residues 1168-1528), rat (Rattus norvegicus, UniProtKB: P41516-1, residues 1168-1526), pig (Sus scrofa, UniProtKB: O46374-1, residues 1171-1533), chicken (Gallus gallus, UniProtKB: O42130-1, residues 1163-1553) and Chinese hamster (Cricetulus griseus (Cricetulus barabensis griseus), UniProtKB: P41515-1, residues 1166-1526) aligned using EMBL-EBI Clustal Omega (Sievers et al, 2011) ordered by input (not alignment). The polybasic motif sequences are highlighted in grey boxes. Bold residues highlight basic residues within the motif sequence (K/R-(Xn=3-7)-K-X-K/R-K/R) of human TOP2A CTD and of corresponding conserved basic residues in other species. The blue box highlights the nucleolar sequence (residues 1192-289) in the rat protein as reported in (Yasuda et al, 2021).

**References**


Sievers, F., Wilm, A., Dineen, D., Gibson, T. J., Karplus, K., Li, W., Lopez, R., McWilliam, H., Remmert, M., Soding, J., Thompson, J. D., & Higgins, D. G. (2011). Fast, scalable generation of high-quality protein multiple sequence alignments using Clustal Omega. Mol Syst Biol, 7, 539. doi:10.1038/msb.2011.75

Yasuda, K., Kato, Y., Ikeda, S., & Kawano, S. (2021). Regulation of catalytic activity and nucleolar localization of rat DNA topoisomerase IIalpha through its C-terminal domain. Genes Genet Syst, 95(6), 291-302. doi:10.1266/ggs.20-00038
